# RNA secondary structure ensemble mapping in a living cell identifies conserved RNA regulatory switches and thermometers

**DOI:** 10.1101/2024.09.16.613214

**Authors:** Ivana Borovská, Chundan Zhang, Edoardo Morandi, Daphne A.L. van den Homberg, Michael T. Wolfinger, Willem A. Velema, Danny Incarnato

**Affiliations:** Department of Molecular Genetics, Groningen Biomolecular Sciences and Biotechnology Institute (GBB), University of Groningen, Groningen, the Netherlands; Institute for Molecules and Materials, Radboud University Nijmegen, Nijmegen, the Netherlands; Department of Theoretical Chemistry, University of Vienna, Vienna, Austria; Research Group Bioinformatics and Computational Biology, Faculty of Computer Science, University of Vienna, Vienna, Austria; RNA Forecast e.U., Vienna, Austria

## Abstract

RNA molecules can populate ensembles of alternative structural conformations, but comprehensively mapping RNA conformational landscapes within living cells presents significant challenges and has, as such, so far remained elusive. Here, we generated the first transcriptome-scale maps of RNA secondary structure ensembles in a living cell, using *Escherichia coli* as a model. Our analysis uncovered features of structurally-dynamic regions, as well as the existence of hundreds of highly-conserved bacterial RNA structural elements. Conditional structure mapping revealed extensive restructuring of RNA ensembles during cold shock, leading to the discovery of several novel RNA thermometers in the 5′ UTRs of the *cspG, cspI, cpxP* and *lpxP* mRNAs. We mechanistically characterized how these thermometers switch structure in response to cold shock and revealed the cspE chaperone-mediated regulation of *lpxP*. Collectively, this work reveals a previously unappreciated complexity of RNA structural dynamics in living cells, and it provides a key resource to significantly accelerate the discovery of regulatory RNA switches.

## Main Text

RNA molecules are pivotal orchestrators of virtually every cellular process, functioning as genetic information carriers and master regulators of gene expression. These roles are intricately intertwined with the ability of RNA molecules to fold into complex structures. Recent strides in the fields of chemical biology and transcriptomics have ushered in a new era, allowing for concurrent examination of the secondary structures of thousands of transcripts in a single experiment^1^. Although the theoretical folding space of RNA molecules is vast^2^, and many RNAs have been reported to populate an ensemble of alternative base-pairing states^3–8^, most transcriptome-wide studies have focused on determining a single conformation for each transcript^9–13^, limiting the predictive power of such models and hindering our understanding of fundamental regulatory mechanisms. To overcome this limitation, a number of methods^3,4,6,7,14–17^, either thermodynamics-dependent or independent^18^, have been devised to deconvolute RNA structural ensembles from chemical probing data. In a recent report we introduced DRACO^4^, an algorithm capable of deconvolving RNA structure ensembles from chemical probing data read out via mutational profiling^14,19,20^ (MaP). In MaP experiments, RNA molecules are first incubated in the presence of a chemical probe, which induces covalent modifications on the RNA at the level of unpaired (or structurally-flexible) nucleotides. Sites of chemical modification are then recorded as cDNA mutations during reverse transcription and decoded via high-throughput sequencing. By analyzing co-mutation patterns in sequencing reads, DRACO can estimate the number of conformations populating the structure ensemble for a given RNA, as well as reconstruct their structures and estimate their relative stoichiometries. Furthermore, unlike the majority of the available ensemble deconvolution methods, which present significant computational overheads, DRACO is optimized for fast computations, hence enabling transcriptome-wide analyses.

In this study, we introduce a novel generalized framework for the identification of functional regulatory RNA structural switches, by combining DRACO-mediated ensemble deconvolution of transcriptome-wide MaP data, and prioritization of functional structures via automated evolutionary conservation assessment, dubbed DeConStruct, and use it to unveil, for the first time, the complexity of the RNA secondary structure ensemble landscape of living *Escherichia coli* cells, on the transcriptome-scale. To achieve this, we subjected exponentially growing *E. coli* DH5α and TOP10 cells to *in vivo* dimethyl sulfate (DMS) probing at 37°C. Ribosomal RNA-depleted samples were subjected to DMS-MaPseq analysis, yielding approximately 1 billion paired-end reads for each experiment. Additionally, we generated libraries from total RNA obtained from both *in vivo* probed cells and *ex vivo* probed RNA following deproteinization, thereby facilitating the derivation of optimized folding parameters (Supplementary Fig. 1 and 2). Analysis of mutation distributions showed an enrichment for mutations on adenine (A) and cytosine (C) bases, constituting on average 56.15 ± 0.55% and 34.15 ± 0.55% of all mutations, respectively (Fig. 1a), as expected from DMS preference for modifying A and C. Area under the Receiver Operator Characteristic (AUROC) curve analysis of 16S and 23S rRNAs confirmed the enrichment of DMS-induced mutations on unpaired bases of rRNA reference structures (AUCs: 0.873-0.914). We observed a remarkable correlation in bulk (ensemble average) DMS reactivities between DH5α and TOP10 cells (*r* = 0.94, Pearson correlation coefficient; Fig. 1b). We achieved a minimum sequencing depth of 5,000X for roughly two-thirds (62-66%) of the bases in the *E. coli* expressed transcriptome (TPM ≥ 10; Fig. 1c), a coverage threshold we had previously shown to be sufficient to ensure robust ensemble deconvolution by DRACO^4^. DRACO-mediated ensemble deconvolution further revealed that, among regions populating an equivalent number of conformations in both strains (Supplementary Table 1), encompassing 1,040,669 bases (accounting for over two-thirds of the analyzed bases), approximately 16.6% populated two or more conformations (Fig. 1d). Importantly, these regions included known riboswitches such as the *lysC* lysine riboswitch^21^, the *ribB* FMN riboswitch^22^, the *thiC* TPP riboswitch^23^, the *mgtL* Mg^2+^ sensor^24^, and the *hisL* histidine leader^25^, as well as the *cspA* RNA thermometer^26^ (Fig. 1e). Under the employed growth conditions, the cognate ligands for these RNA switches are expected to be abundant. Accordingly, the Mg^2+^ sensor predominantly adopted the conformation encompassing stem loops A and B (conformation B: 65.15 ± 0.45%; Fig. 1e, top inset), which is favored at high Mg^2+^ concentrations. Similarly, the histidine leader predominantly favored the attenuated conformation (conformation A: 70.4 ± 0.4%; Fig. 1e, bottom inset), which facilitates complete leader peptide translation in the presence of high histidyl-tRNA levels. Altogether, these results confirm that our approach is indeed suited for the identification of regulatory RNA structural switches from transcriptome-wide chemical probing data.

**Fig. 1.**
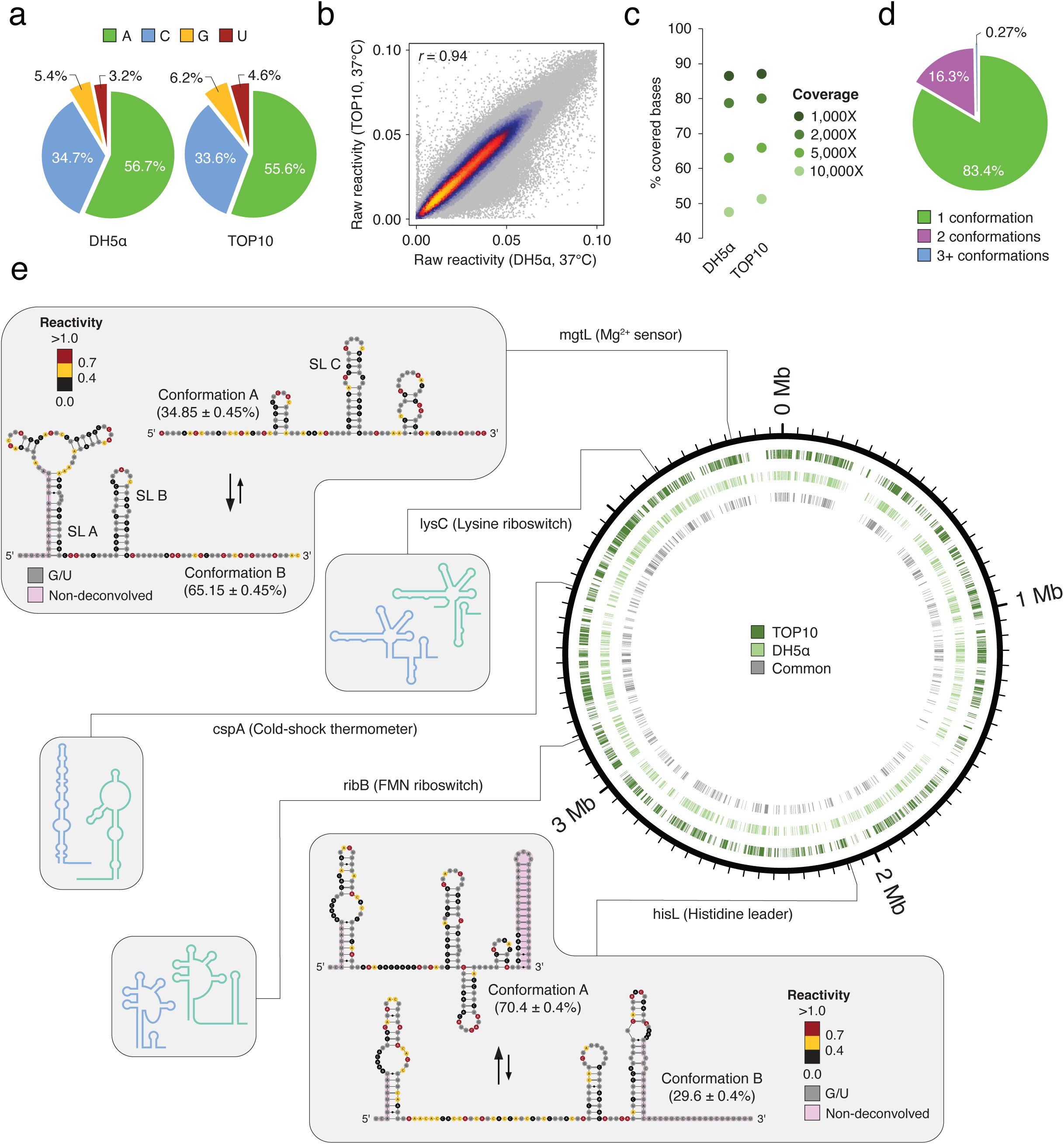
*In vivo* mapping of *Escherichia coli* RNA structure ensembles. (**a**) Pie charts depicting the percentages of mutated bases across the transcriptomes of DH5α and TOP10 cells upon DMS probing at 37°C. (**b**) Heat scatter-plot of raw DMS reactivities across bases with coverage ≥ 10,000X in the transcriptome of DH5α and TOP10 cells. Outliers (bases with mutation frequency > 0.1) were excluded. (**c**) Percentage of bases covered in the expressed transcriptome (TPM ≥ 10) of DH5α and TOP10 cells, at different sequencing depths. (**d**) Pie-chart depicting the percentages of bases in the *E. coli* transcriptome populating 1, 2, or 3+ conformations. Only bases populating the same number of conformations in DH5α and TOP10 cells were considered. (**e**) Schematic representation of the *E. coli* genome. Regions populating 2+ conformations in TOP10 (dark green), DH5α (light green), or both (grey) are indicated. Examples of riboswitches and RNA thermometers known to populate 2 alternative conformations are shown. For the *mgtL* Mg^2+^ sensor and the *hisL* histidine leader, reconstructed reactivities for the two conformations have been averaged across DH5α and TOP10 cells and overlaid on the known structures. Bases falling outside of the region deconvolved by DRACO are marked in pink.

To elucidate the characteristics distinguishing regions populating a single conformation (hereafter referred to as “1 regions”) from those exhibiting two or more conformations (hereafter referred to as “2+ regions”), we first examined the possibility that the separation between these regions might result from differences in their information content. As DRACO relies on A/C co-mutation patterns to perform ensemble deconvolution, we wondered whether 2+ regions might be enriched in A/C bases and thus might possess higher information content, but no such enrichment was observed (AC% 1 regions: 49.6%, 2+ regions: 49.1%, p-value: 0.99, one-tailed Wilcoxon rank sum test). Subsequently, we performed partition function folding, either unconstrained or constrained by bulk DMS-MaPseq reactivities and optimized folding parameters (see Materials and Methods; Supplementary Fig. 2), thereby deriving base-pairing probabilities across the entire *E. coli* transcriptome. We calculated median Shannon entropies, a measure of the structural disorder across each base of a transcript, for both 1 and 2+ regions. Unexpectedly, Shannon entropies were significantly higher for 1 regions than for 2+ regions in both unconstrained (p-value: 2.8e-12, Wilcoxon rank sum test; Fig. 2a) and experimentally-constrained predictions (p-value: 5.1e-28, Wilcoxon rank sum test; Supplementary Fig. 3a). This result suggested that 2+ regions may predominantly occupy well-defined structural states, while 1 regions may exhibit greater disorder and exist in numerous states with lower probability. In line with this hypothesis, we observed that the median probability of bases to be unpaired was significantly higher in 1 regions compared to 2+ regions in unconstrained (p-value: 2.8e-39, Wilcoxon rank sum test; Fig. 2b) and even more prominently in experimentally-constrained (p-value: 2.0e-103, Wilcoxon rank sum test; Supplementary Fig. 3b) predictions. Accordingly, median DMS reactivities were significantly higher, and Gini indexes significantly lower, in 1 regions as compared to 2+ regions, indicating that 1 regions tend to be less structured than 2+ regions (median reactivity p-value: 5.9e-89, Gini index p-value: 1.2e-109, Wilcoxon rank sum test; Fig. 2c, d). The propensity of 2+ regions towards increased structural orderliness appeared to be largely sequence-driven, as these regions exhibited a significantly higher GC% content (p-value: 7.4e-15), as well as lower folding free energies than expected for sequences of same dinucleotide composition (p-value: 1.3e-27, Wilcoxon rank sum test; Supplementary Fig. 3c), as compared to 1 regions. Furthermore, comparative sequence analysis of 10 Gram-negative genomes revealed that 2+ regions are significantly more conserved than 1 regions (p-value: 2.8e-57, Wilcoxon rank sum test; Fig. 2e).

**Fig. 2.**
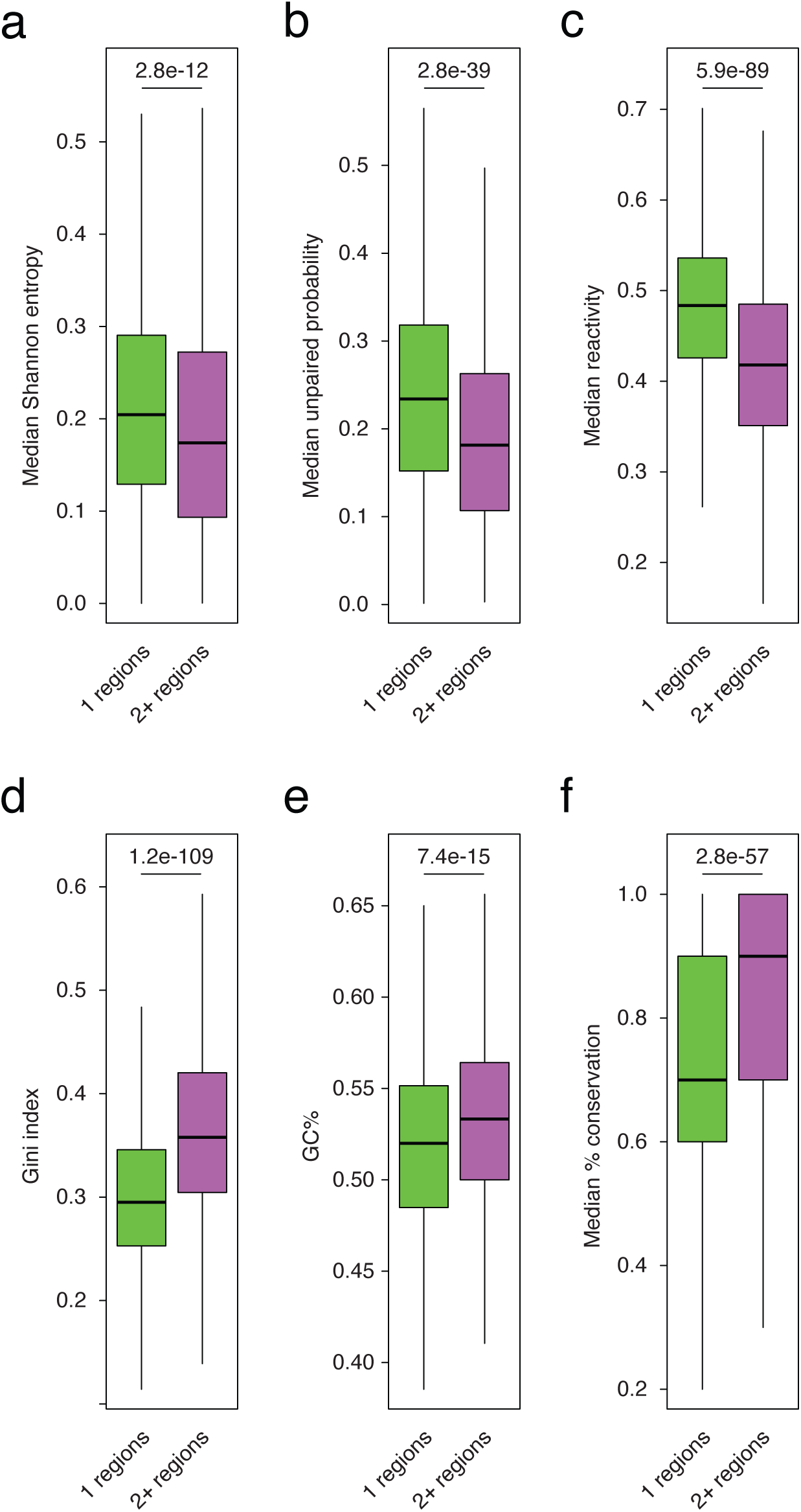
Features of structurally-heterogeneous RNA regions. Box-plots depicting the distributions for different features across regions populating 1 or 2+ conformations *in vivo*. (**a**) Median Shannon entropies (from unconstrained predictions). (**b**) Median unpaired probabilities (from unconstrained predictions). (**c**) Median bulk (ensemble average) *in vivo* DMS reactivities. (**d**) Gini indexes calculated on bulk (ensemble average) *in vivo* DMS reactivities. (**e**) GC% content. (**f**) Median % sequence conservation calculated on a set of 10 Gram-negative bacterial genomes. For all plots, boxes span the 25^th^ to the 75^th^ percentile. The center represents the median. Outliers (values below the 25^th^ percentile – 1.5 times the IQR, or above the 75^th^ percentile + 1.5 times the IQR) are not shown. P-values are calculated using the Wilcoxon rank sum test.

We next wondered whether the observed structural heterogeneity might arise from alternative transcript isoforms generated by alternative promoters and/or terminators^27^. We tested whether 2+ regions were enriched within transcripts generated from alternative promoters but did not observe any enrichment (expected: 10.4%, observed: 9.7%, p-value = 0.75, one-tailed binomial test). Similarly, no enrichment was observed for genes harboring alternative terminators (expected: 4.26%, observed: 3.86%, p-value = 0.73, one-tailed binomial test). Furthermore, we ruled out the possibility that structural heterogeneity might preferentially arise as a consequence of interactions with small RNAs (sRNAs), as no significant enrichment was observed for experimentally-validated sRNA interactions^28^ (p-value: 0.86, one-tailed binomial test). We next investigated two additional potential confounding factors: RNA translation and RNA decay. To evaluate the propensity of 2+ regions towards higher structural heterogeneity in absence of any contribution by cellular factors, we extracted RNA from exponentially growing DH5α and TOP10 cells and subjected it to *in vitro* refolding and DMS-MaPseq analysis. We achieved high correlations and coverage as we did for the in-cell datasets (Supplementary Fig. 4a, b). Overall, *in vitro* refolded RNAs showed a higher fraction of structurally-heterogeneous regions (∼32.3%, Supplementary Fig. 4c and Supplementary Table 2) as compared to *in vivo* samples. These regions included known RNA switches such as the Mg^2+^ sensor, the molybdenum cofactor (Moco) riboswitch^29^, and the *cspA* RNA thermometer (Supplementary Fig. 4d-f). Notably, we observed that over two-thirds (67.3%) of the *in vivo*-identified 2+ regions appeared to be structurally-heterogeneous under *in vitro* conditions as well, with 94.1% of them populating the same number of conformations in cell and *in vitro*, and 5.9% showing increased structural heterogeneity under *in vitro* conditions (Supplementary Fig. 4G and Supplementary Table 3). It is also worth noticing that, upon reanalysis of published ribosome profiling data^30^ we observed that, regions exhibiting structural heterogeneity under *in vitro* conditions but not in cell, showed significantly higher ribosome occupancy and translation efficiencies as compared to regions being heterogeneous both *in vivo* and *in vitro*, or only *in vivo* (p-values: 3.4e-8 and 9.3e-10, Wilcoxon rank sum test; Supplementary Fig. 4h). This is in line with previous findings^13,31^ showing that translation can actively unfold RNA structures in cells, hence suggesting that it might be partly masking the actual structural heterogeneity of cellular RNAs.

We further polished this high-confidence set of 2+ regions by discarding those overlapping with experimentally-determined RNase E cleavage sites^32^, for which the observed heterogeneity might derive from the presence of decay fragments. Importantly, feature reanalysis for this subset did not affect the aforementioned differences between 1 and 2+ regions (Supplementary Fig. 5). Furthermore, as 1 regions might be polluted by putative 2+ regions that were not deconvolved by DRACO, we also analyzed the same set of features in high-confidence 2+ regions as compared to a random set of transcriptome regions of matching size and observed the exact same trends (Supplementary Fig. 6). Interestingly, the high-confidence subset showed an even higher median % conservation with respect to both 1 regions and random transcriptome regions (Supplementary Fig. 5e and Supplementary Fig. 6e), indicating that 2+ regions might be enriched for conserved RNA structural regulatory elements.

To evaluate this possibility, we first identified a total of 901 structurally-heterogeneous regions whose DRACO-deconvolved reactivity profiles could be non-ambiguously matched between DH5α and TOP10 cells *in vivo* data and generated experimentally-informed RNA structure models. We next adopted the DeConStruct framework, which builds on top of the *cm-builder* method we previously introduced^33,34^, but significantly expanded to automatically build alignments of related sequences from a representative set of bacterial genomes, and validated it on known RNA switches (Supplementary Fig. 7). After discarding regions encompassing any known RNA structure element from RFAM and applying stringent alignment selection criteria (see Materials and Methods), we identified 226 regions (∼26.5%) for which at least one of the conformations showed robust covariation support, generally regarded as a strong evidence of RNA structure functionality, as determined by R-scape ^35^ analysis. Taken together, our data hints at the existence of a previously unappreciated repertoire of functional regulatory RNA elements, likely including novel regulatory RNA switches, in bacterial RNAs. To facilitate the analysis and exploration of these regions, we further aggregated them into a browsable web-site (see Data and materials availability). It is worth emphasizing that, due to a number of different factors (namely, errors in the thermodynamic model and in the resulting structure predictions, the employed set of representative bacterial genomes, and the stringent alignment selection criteria), this set likely represents an underestimate of the actual number of conserved, structurally-heterogeneous regions in the *E. coli* transcriptome. Accordingly, approximately 40% of the regions showed at least 1 covarying base-pair.

We next investigated how RNA secondary structure ensembles get redistributed in response to environmental cues. We chose cold shock as changes in RNA structure have been previously reported to be one of the hallmarks of cold adaptation in bacteria^30,36^. We therefore performed DMS-MaPseq analysis of exponentially growing *E. coli* cells shocked at 10°C for 20 minutes. It has been previously shown that DMS reaction kinetics is slower at 10°C than it is at 37°C and, therefore, reaction times need to be increased to achieve similar modification rates^11,30^. We wondered whether this might cause artifacts as, during the timeframe of DMS modification, which would exceed the average half-life of *E. coli* mRNAs upon cold shock^37^, the expressed transcriptome changes substantially. To this end, we shocked *E. coli* cells for 20 minutes at 10°C, treated them with DMS for 2 or 30 minutes, and compared their expression profiles to those of DMS-untreated cells by RNA-seq (Supplementary Fig. 8a). Gene expression analysis showed that, while DMS-untreated cells underwent the expected transcriptome changes between 2 and 30 minutes (e.g., a robust upregulation of mRNAs encoding for cold-induced cold shock proteins, CSPs) (R^2^ = 0.88, Pearson correlation coefficient), no change occurred for cells treated with DMS (R^2^ = 0.99, Pearson correlation coefficient). This data indicates that, despite slowed down reaction kinetics at 10°C, addition of DMS nearly immediately blocks all cellular processes, including transcription and RNA decay, hence providing an instantaneous snapshot of the RNA structurome.

Cold shocked cells showed extremely well correlated DMS reactivities (*r* = 0.95, Pearson correlation coefficient; Supplementary Fig. 8b), and formed a well-separated cluster with respect to exponentially growing cells at 37°C in PCA analysis (Supplementary Fig. 8c), indicating the existence of substantial structural rearrangements at 10°C. Surprisingly, DRACO-mediated ensemble deconvolution showed that, among regions populating an equivalent number of conformations in both strains (Supplementary Table 4), totaling 807,853 bases, approximately 32.6% populated two or more conformations, corresponding to a nearly twofold increase as compared to cells grown at 37°C (Fig. 3a). When focusing solely on regions covered both at 37°C and 10°C, encompassing 556,645 bases, we observed that over 5 times more regions showed increased ensemble heterogeneity, and thus populated a higher number of conformations, at 10°C, than those with reduced ensemble heterogeneity (increased heterogeneity at 10°C: ∼15.7%, decreased heterogeneity at 10°C: ∼3%; Fig. 3b and Supplementary Table 5). While this increase in structural diversity upon cold shock might seem counterintuitive to traditional thermodynamic expectations, which predict fewer structural states at lower temperatures, we suggest that the decrease in temperature might reduce entropy of 1 regions, promoting higher structuredness and fewer, more well-defined, structural states.

**Fig. 3.**
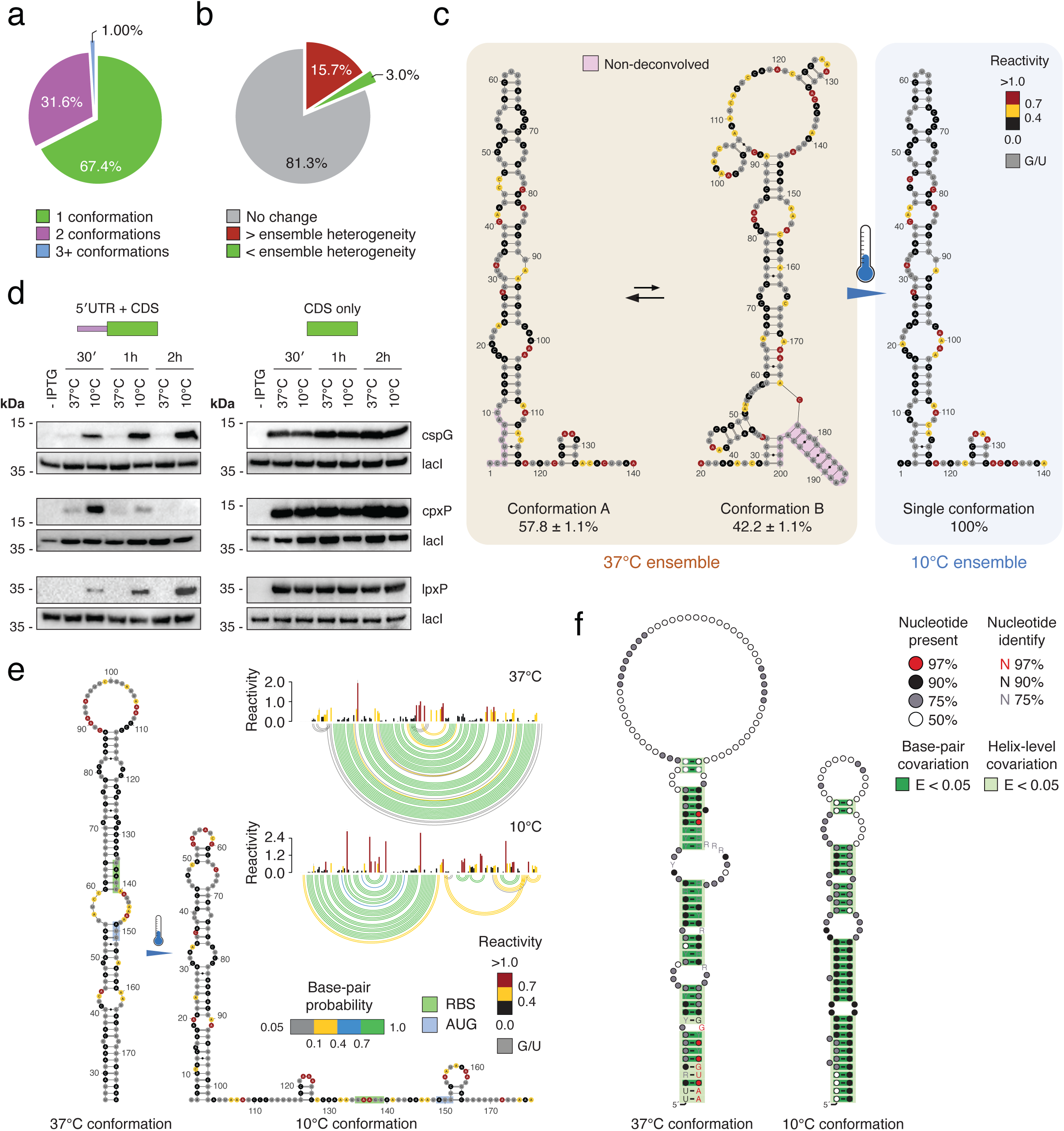
Redistribution of RNA structure ensembles upon cold shock. (**a**) Pie-chart depicting the percentages of bases in the *E. coli* transcriptome populating 1, 2, or 3+ conformations after cold shock. Only bases populating the same number of conformations in DH5α and TOP10 cells are considered. (**b**) Pie-chart depicting the percentages of bases in the *E. coli* transcriptome, for transcripts expressed both at 37°C and at 10°C, for which the ensemble heterogeneity increases (red), decreases (green), or remains unchanged (grey) after cold shock. (**c**) RNA secondary structure ensemble analysis of the *cspA* RNA thermometer at 37°C (left), and after cold shock (right). Reactivities are averaged across DH5α and TOP10 cells and overlaid on the predicted structures. Bases falling outside of the region deconvolved by DRACO are marked in pink. (**d**) Western blot analysis of cspG, cpxP and lpxP expression at 37°C and 10°C, 30 minutes, 1 hour and 2 hours post-IPTG induction and cold shock for constructs harboring both the 5′ UTR and the CDS (left), or only the CDS (right). lacI is the loading control. (**e**) Secondary structure models of the 5′ UTR of *cspG* at 37°C and 10°C, with overlaid *in vitro* DMS reactivities, along with reactivity profiles and base-pairing probabilities for both conformations. Reactivities are averaged across two independent experiments. Error bars represent the standard deviation. (**f**) Structure models for the two conformations of the identified *cspG* RNA thermometer, inferred by phylogenetic analysis. Base-pairs showing significant covariation (as determined by R-scape) are boxed in dark green (E-value < 0.05). Helices showing helix-level covariation support (E-value < 0.05) are boxed in light green.

Among the fewer regions that exhibited reduced ensemble heterogeneity upon cold shock, we observed the well-known *cspA* RNA thermometer^26^. Previous studies proposed that *cspA* can switch between a translationally-incompetent conformation at 37°C and a translationally-competent conformation at 10°C^26,30^. However, such a model cannot explain why, at 37°C, cspA is one of the top 10% expressed proteins in the *E. coli* proteome^38^. Accordingly, ensemble deconvolution analysis showed that, at 37°C, the 5′ UTR of *cspA* populates two conformations, with the translation-competent conformation being the predominant one (conformation A: 57.8 ± 1.1%) and becoming the sole conformation upon cold shock (Fig. 3c). Interestingly, the two conformations of *cspA* could also be observed in our *in vitro* refolded dataset (Supplementary Fig. 5e), albeit with inverted stoichiometries (conformation A: 45.25 ± 0.55%; conformation B: 54.75 ± 0.55%), hence indicating that the cellular environment plays a key role in determining conformation abundances in the ensemble. Ontology analysis of genes containing regions undergoing ensemble redistribution upon cold shock showed a significant enrichment for terms associated with response to temperature changes (Response to heat: 2.5e-6, Response to cold: 3.8e-5, Cellular response to heat: 2.0e-2), stress response (p-value: 6.2e-3), and pathways commonly modulated in response to cold shock, such as glycolysis (p-value: 1.4e-6), fatty acid biosynthesis (p-value: 1.5e-4), lipid biosynthesis and lipid metabolism (p-value: 8.0e-6 and 1.1e-3), as well as protein folding and unfolding (p-values: 2.5e-3 and 9.5e-5), among others. Additionally, reanalysis of publicly available ribosome profiling data^30^ indicated a moderate yet significant increase in translation efficiency for both genes encompassing regions of increased or decreased structural heterogeneity upon cold shock, 10 minutes after the temperature shift to 10°C, as compared to 37°C (p-value: 7.9e-7, paired Wilcoxon rank sum test; Supplementary Fig. 8d). This increase was not observed for genes encompassing regions whose RNA ensemble heterogeneity remained unchanged. Collectively, these data suggest the existence of a wide catalog of undiscovered RNA thermometers in bacteria.

To further explore this possibility, we selected three genes whose 5′ UTRs encompassed regions of increased structural heterogeneity upon cold shock, and that displayed increased translation efficiency at 10°C by ribosome profiling, as well as structural conservation, namely *cspG*, *cpxP*, and *lpxP*. The *cspG* mRNA encodes for a cold shock protein, which belongs to the same family of cspA. *cspG* has been previously shown to be robustly induced by cold shock at the transcriptional level^39,40^ but, to the best of our knowledge, its regulation at the translational level, as well as the role of its 5′ UTR as RNA thermometer, have never been reported. The *cpxP* mRNA encodes for a periplasmic protein involved in sensing and mediating the adaptation to various cellular stresses that might result in protein misfolding^41,42^, which is crucial for heat as well as cold shock responses^43^. The *lpxP* mRNA encodes for a palmitoleoyl transferase that catalyzes the palmitoylation of lipid A to maintain optimal outer membrane fluidity at low temperature^44^. We first tested whether the 5′ UTRs of these mRNAs played a role in regulating their translation upon cold shock by cloning these genes, with or without their 5′ UTRs, in IPTG-inducible constructs, and by measuring their expression at 37°C versus 10°C. All three genes showed minimal to no expression at 37°C, but robust translation upon cold shock, while deletion of their 5′ UTRs abrogated their cold-mediated regulation (Fig. 3d). *cspG* and *lpxP* both showed increasing translation over a 2-hour time course, while *cpxP* expression quickly increased within the first 30 minutes of cold shock, and then rapidly decreased after 1 hour.

We next wondered whether temperature shift alone would be sufficient to remodel the structure of the 5′ UTR of these genes. We therefore performed *in vitro* transcription of these mRNAs, either at 37°C or 10°C, followed by DMS-MaPseq analysis. Notably, both *cspG* and *cpxP* showed substantial structural rearrangements at 10°C as compared to 37°C (Fig. 3e and Supplementary Fig. 9a). These novel temperature-induced RNA structural switches were supported by extensive covariation (Fig. 3f and Supplementary Fig. 9b), underscoring their functional relevance. We further asked whether the regulation observed for the *cspG* 5′ UTR was also shared by other cold-induced members of the csp family. Particularly, *cspB*’s 5′ UTR shares extensive sequence identity with the 5′ UTR of *cspG* (Supplementary Fig. 10a), while that of *cspI* diverges. *cspI* has been previously shown to be subjected to translational control by its 5′ UTR but, just like for *cspG*, its putative role as an RNA thermometer has never been investigated^45^. Intriguingly, all three 5′ UTRs showed a similar behavior, characterized by a large stem-loop structure encompassing the entire 5′ UTR, as well as part of the coding region, at 37°C, which sequestered both the ribosome binding site (RBS) and the start codon, and a register-shifted, shorter, stem-loop structure involving the sole 5′ UTR at 10°C, which left the RBS and the start codon available for translation initiation (Fig. 3e and Supplementary Fig. 10b, c).

Analogously to the 5′ UTRs of csp-encoding mRNAs, the 5′ UTR of *lpxP* showed two nearly equimolar conformations at 10°C in cell (Fig. 4a), respectively characterized by a stem-loop structure (SL, conformation A) sequestering both RBS and start codon, which is also the predominant conformation at 37°C (Fig. 4b), and a register-shifted stem-loop (SL_alt_, conformation B) leaving both elements available for translation initiation. Both SL and SL_alt_ showed extensive covariation support (Fig. 4c), and covariance model-guided homology search revealed the existence of homologous structures in several other Gram-negative bacteria (Supplementary Fig. 11). We confirmed that conformation B corresponded to the translation-competent conformation by generating an SL_alt_-stabilized mutant and validated its structure by targeted DMS-MaPseq analysis (Fig. 4d). Stabilization of SL_alt_ abrogated the cold-mediated regulation of *lpxP*, leading to its constitutive expression at both 37°C and 10°C (Fig. 4e). To further confirm that the observed regulation was indeed structure-mediated, rather than being caused by other factors such as, for example, altered mRNA decay of the SL_alt_-stabilized mutant, we adopted the PURE system^46^, a reconstituted *E. coli in vitro* transcription-translation system. Expression of *lpxP* harboring the wild type 5′ UTR could not be detected after 2 hours at 37°C, while both the CDS-only template and the SL_alt_-stabilized mutant were robustly translated (Fig. 4f, g).

**Fig. 4.**
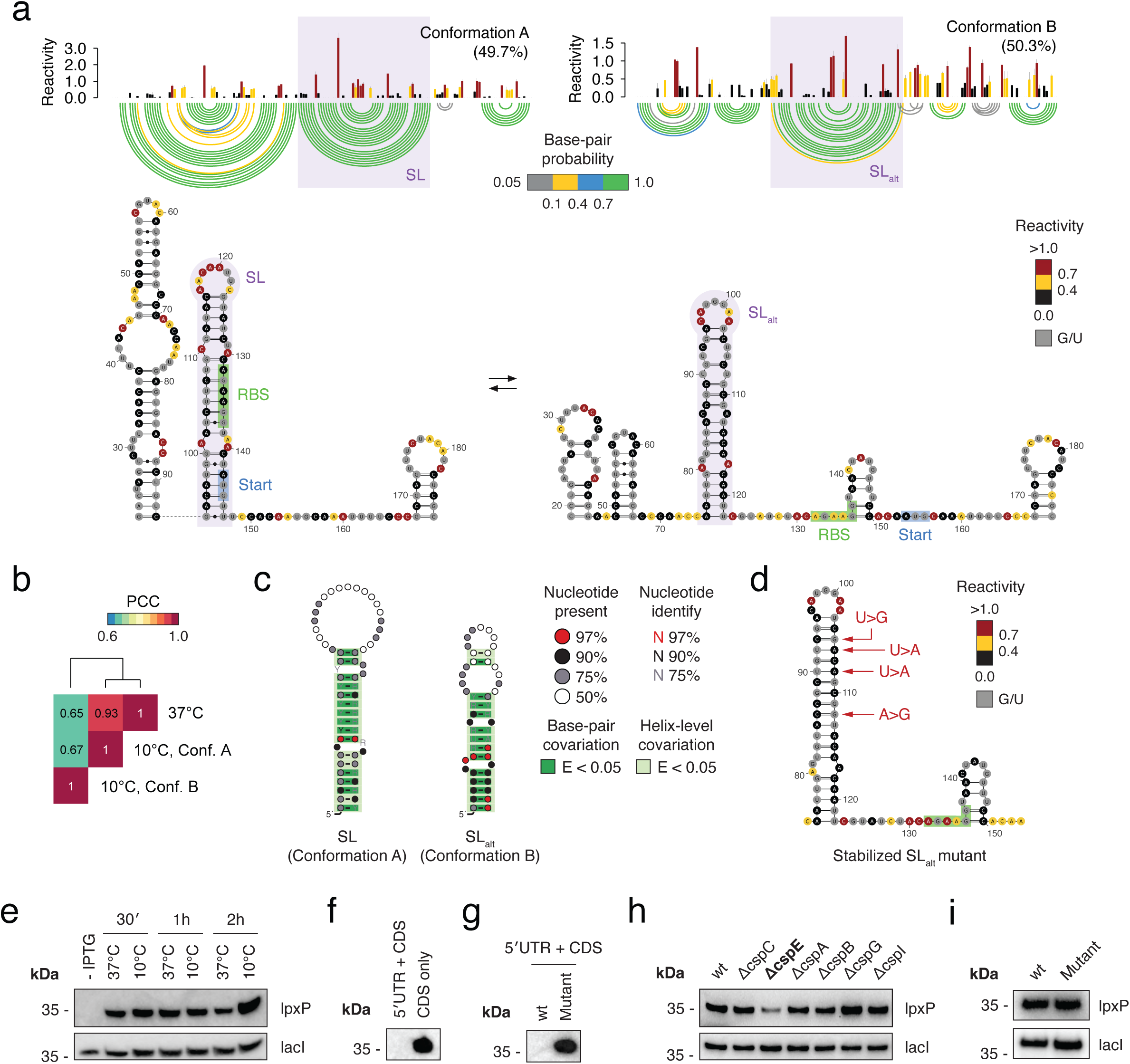
Characterization of the *lpxP* RNA thermometer. (**a**) Secondary structure models for the two conformations of the *lpxP* 5′ UTR as identified via ensemble deconvolution analysis of cold shocked bacteria, with overlaid *in vivo* DMS reactivities at 10°C, along with reactivity profiles and base-pairing probabilities for both conformations. Reactivities are averaged across DH5α and TOP10 cells. Error bars represent the standard deviation. The two register-shifted stem-loops SL and SL_alt_ are highlighted in purple. (**b**) Heatmap of pairwise Pearson correlation coefficients (PCC) of normalized DMS reactivities across the two alternative conformations of the *lpxP* 5′ UTR at 10°C, and the 37°C conformation. (**c**) Structure models for the SL and SL_alt_ stem-loops, inferred by phylogenetic analysis. Base-pairs showing significant covariation (as determined by R-scape) are boxed in dark green (E-value < 0.05). Helices showing helix-level covariation support (E-value < 0.05) are boxed in light green. (**d**) Secondary structure of the SL_alt_-stabilized mutant, with overlaid DMS reactivities. (**e**) Western blot analysis of SL_alt_-stabilized mutant expression at 37°C and 10°C, 30 minutes, 1 hour and 2 hours post-IPTG induction and cold shock. lacI is the loading control. (**f**) Western blot analysis of the full-length (5′ UTR + CDS), or CDS-only *lpxP* constructs, translated using the PURE system. (**g**) Western blot analysis of the full-length (5′ UTR + CDS) wild type and SL_alt_-stabilized mutant constructs, translated using the PURE system. (**h**) Western blot analysis of full-length (5′ UTR + CDS) *lpxP* expression at 10°C, 1-hour post-IPTG induction, in wild type or in CSP single knock out clones from the KEIO collection. lacI is the loading control. (**I**) Western blot analysis of SL_alt_-stabilized mutant expression at 10°C, 1-hour post-IPTG induction, in wild type or in cspE knock out bacteria.

Unlike the 5′ UTRs of csp-encoding mRNAs and of *cpxP*, however, *lpxP*’s 5′ UTR showed no structure rearrangement between 37°C and 10°C under *in vitro* conditions, solely populating the translation-incompetent conformation at both temperatures (Supplementary Fig. 12). We hypothesized that the energy barrier between the translation-competent and translation-incompetent conformations might be so high that switch from SL to SL_alt_ could not happen spontaneously on a biologically-relevant time scale, and that, as such, it might be a chaperone-assisted process. Interestingly, reanalysis of published cspC and cspE CLIP-seq data from septicemic *E. coli*^47^ showed binding of both proteins to the 5′ UTR of *lpxP*. Similarly, RIP-seq analysis of csp proteins in *Salmonella enterica* serovar Typhimurium^48^, which we found to also carry a homologous structural switch (Supplementary Fig. 11), showed binding of both cspE and, to a lesser extent, cspC to *lpxP*’s 5′ UTR, hinting at a highly conserved regulatory mechanism. To confirm the role of csp proteins in regulating the structural switch in *lpxP*’s 5′ UTR we analyzed *lpxP* translation in *E. coli* csp knock outs from the KEIO collection^49^, and observed a ∼50% reduction in lpxP expression in ΔcspE cells (Fig. 4h), which was not observed for the SL_alt_-stabilized mutant (Fig. 4i).

In summary, we reported here the first transcriptome-scale map of RNA secondary structure ensembles in a living cell, and introduced a generalized framework, DeConStruct, that, by combining chemical probing-guided ensemble deconvolution and analysis of evolutionary conservation by covariation, accelerates the discovery of novel, functional regulatory RNA structural switches that have remained so far largely elusive. By leveraging this framework, we report the discovery of hundreds of candidate conserved RNA structural switches, which will provide an important resource for the identification of novel riboswitch classes, as well as novel RNA thermometers. We further experimentally characterize them, demonstrating that our approach can recover both canonical, protein-independent thermometers, such as *cspG*, *cspI* and *cpxP*, as well as chaperone-dependent ones, such as *lpxP*. Furthermore, as the *lpxP* thermometer represents a true on-off temperature-controlled switch, oppositely to *cspA*, we can anticipate it will have important applications in synthetic biology.

Our data further challenges the traditional view of the cold shock response in bacteria, which has long been understood primarily in terms of RNA unfolding facilitated by the overexpression of cold shock proteins, or the increased structural rigidity of RNA due to lower temperatures. Instead, our findings suggest that the response is far more intricate, involving a complex redistribution of RNA structural ensembles. This indicates that cold shock induces a broader and more dynamic reorganization of RNA structures than previously thought, reflecting a sophisticated cellular adaptation to temperature changes.

Albeit representing a crucial step towards a better understanding of the regulatory roles of RNA structures in living cells, as well as the nuanced dynamics of RNA structure ensembles in response to environmental cues, the current study presents a number of limitations. Firstly, the use of DMS limits chemical probing to the interrogation of A and C bases, which might hamper the identification of small and A/C-poor structurally-dynamic regions. This will hopefully be addressed in future studies, by taking advantage of recent advances in chemical probing protocols and reagents^50,51^, which allow querying all four nucleotides. Secondly, an implicit assumption of chemical probing-guided ensemble deconvolution analyses is that, in order to identify coexisting alternative structural states for an RNA, they need to interconvert at a rate that is slower than the timescale of the probing experiment. Indeed, transition barrier analysis for the alternative conformations identified in this study showed that, in general, they tend to be separated by high energy barriers (median at 37°C: 8.6 kcal/mol, median at 10°C: 13.5 kcal/mol; Supplementary Fig. 13a), and that their interconversion requires the disruption/creation of, on average, at least 50% of the base-pairs (52-57%; Supplementary Fig. 13b), hence suggesting that they are unlikely to spontaneously interconvert on a biologically relevant timescale. Rather, their interconversion in the cell is likely a chaperone-mediated process. Accordingly, 2+ regions tend to be significantly enriched for binding of the chaperones cspC and cspE as compared to 1 regions (cspC p-value: 2.2e-28, cspE p-value: 3.9e-39, one-tailed binomial test). Oppositely, as previously discussed, it is possible that “1 regions” might represent an average of short-lived (i.e., excited) states, interconverting at a faster rate, or that they might sample alternative conformations at stoichiometries too low (<5-10%) to be detected by chemical probing-guided ensemble deconvolution analyses.

Nevertheless, although the structural switches identified in this study might only represent a conservative estimate of the actual RNA structural diversity in *E. coli*, our findings hold great significance both to advance our understanding of gene expression regulation, as well as for the development of innovative antimicrobial RNA-targeted therapeutic strategies^52^.

## Acknowledgments

Knock out clones from the KEIO collection were kindly provided by Dr. Silke R. Bonsing-Vedelaar (University of Groningen) and Prof. Matthias Heinemann (University of Groningen). We would also like to thank Dr. Anton I. Petrov (Riboscope Ltd.), Dr. Steve L. Bonilla (The Rockefeller University) and Dr. Alisha Jones (New York University) for their helpful comments.

## Author contributions

Conceptualization: IB, DI

Wet-lab: IB, CZ, DAL, WAV, DI

DRACO algorithm development: EM, DI

Bioinformatics, structure modeling, data analysis: EM, MTW, DI

Writing: IB, EM, CZ, MTW, DAL, WAV, DI

## Data availability

Sequencing data have been deposited to the Gene Expression Omnibus (GEO) database, under the accession GSE247244. Raw MM files for analysis with DRACO are available from Zenodo (https://doi.org/10.5281/zenodo.10357457). Additional processed files (including the browsable set of conserved structurally-heterogeneous regions) are available at https://www.incarnatolab.com/datasets/EcoliEnsembles_Borovska_2024.php.

The source codes of DRACO v1.2, of the *cm-builder*, *filterMM*, *consensusFold* and *stockholmPolish* utilities, and the *DeConStruct* pipeline are freely available from GitHub, under the GPLv3 license (https://github.com/dincarnato/draco, https://github.com/dincarnato/labtools, and https://github.com/dincarnato/papers).

