## Supplementary Materials for "RNA secondary structure ensemble mapping in a living cell identifies conserved RNA regulatory switches and thermometers"

### Methods

<u>Strains, growth conditions and *in vivo* DMS probing.</u>

E. coli K-12 MG1655 derivative strains DH5α and TOP10 were streaked on LB plates, and a single colony was picked, inoculated in 4 mL LB broth, and grown overnight at 37°C with shaking. The day after, the culture was diluted to an $OD_{600}$ = 0.05 in 25 mL LB broth and grown at 37°C until $OD_{600} \approx 0.5$ (~2 h). For cold shock, 2 mL of this culture were mixed with 2 mL of LB broth pre-chilled to 0°C in a water-ice slurry, then incubated at 10°C for 20 min. For dimethyl sulfate (DMS; cat. D186309, Merck) probing, DMS from a fresh 1:4 dilution in ethanol (~2.64 M) was added to the bacteria at a final concentration of 200 mM. Probing was conducted for 2 min at7°C, or for 30 min at 10°C (to achieve comparable modification efficiencies), with moderate shaking (800 RPM). Reactions were then quenched by addition of 1 volume 1 M DTT, after which bacteria were collected by centrifugation at 17,000g for 1 min. Supernatant was discarded, the pellet was washed twice with 0.5 M DTT, and then immediately subjected to RNA extraction.

<u>RNA extraction.</u>

Cell pellets were resuspended in 62.5 µl Resuspension Buffer [20 mM Tris-HCl pH 8.0; 80 mM NaCl; 10 mM EDTA pH 8.0], supplemented with 100 µg/mL final Lysozyme (cat. L6876, Merck) and 20 U SUPERase•In™ RNase Inhibitor (cat. A2696, ThermoFisher Scientific), by vigorous vortexing. Samples were incubated at room temperature for 1 min, followed by addition of 62.5 µl Lysis Buffer [0.5% Tween-20; 0.4% Sodium deoxycholate; 2 M NaCl; 10 mM EDTA]. Samples were then inverted 5-10 times and incubated at room temperature for 2 min, followed by additional 2 min on ice. 1 mL ice-cold TRIzol™ Reagent (cat. 15596018, ThermoFisher Scientific) was then added, samples were vigorously vortexed for 15 sec, and RNA was extracted as per manufacturer instructions. Residual gDNA was removed by digestion with TURBO DNase I (cat. AM2239, ThermoFisher Scientific) at 37°C for 30 min.

<u>DMS probing of *in vitro* refolded RNA.</u>

10 µg of total RNA from exponentially growing *E. coli* were diluted in 89 µl nuclease-free water, then heat denatured at 95°C for 2 min and immediately chilled on ice for 1 min. 10 µl of ice-cold 10X Folding Buffer [250 mM HEPES pH 7.5; 2 M KCl] were then added and samples were incubated at 37°C for 15 min. 1 µl 500 mM $MgCl_2$ (pre-warmed at 37°C) was then added and samples were incubated at 37°C for additional 15 min to enable tertiary structure formation. Probing was conducted by adding DMS at a final concentration of 200 mM and incubating the samples at 37°C for 2 min. Reactions were then quenched by addition of 1 volume 1 M DTT, after which RNA was cleaned up on Monarch® RNA Cleanup columns (cat. T2030L, New England Biolabs) as per manufacturer instructions.

<u>Extraction and DMS probing of native deproteinized rRNA.</u>

Native deproteinized *E. coli* rRNA was prepared as previously described[33]. Briefly, 2 mL of DH5α or TOP10 cells grown to an $OD_{600} \approx 0.5$ were collected by centrifugation at 1000g for 5 min (4°C), then resuspended in 1 mL Resuspension Buffer [15 mM Tris–HCl pH 8.0; 450 mM Sucrose;

8 mM EDTA], supplemented with 100 µg/mL final Lysozyme. Samples were incubated at 22°C for 5 min, then on ice for additional 10 min, after which protoplasts were collected by centrifugation at 5000g for 5 min (4°C). Protoplast pellet was then resuspended in 120 µl Protoplast Lysis Buffer [50 mM HEPES pH 8.0; 200 mM NaCl; 5 mM MgCl2; 1.5% SDS], supplemented with 0.2 µg/µl Proteinase K (cat. P2308, Merck), and samples were incubated at 22°C for 5 min, followed by 5 min on ice. SDS was precipitated by addition of 30 µl SDS Precipitation Buffer [50 mM HEPES pH 8.0; 1 M Potassium acetate; 5 mM MgCl2], followed by centrifugation at 17000g for 5 min (4°C). Supernatant was extracted twice with phenol:chloroform:isoamyl alcohol (25:24:1), pre-equilibrated in RNA Folding Buffer [50 mM HEPES pH 8.0; 200 mM NaCl; 5 mM MgCl2], and twice with chloroform. Deproteinized samples were then supplemented with 20 U SUPERase•In™ RNase Inhibitor equilibrated at 37°C for 20 min. DMS from a 1:4 dilution in ethanol was added to a final concentration of 200 mM and samples incubated at 37°C for 2 min with shaking (800 RPM). Reactions were quenched by addition of 1 volume 1 M DTT, then cleaned up using Monarch® RNA Cleanup columns as per manufacturer instructions.

DMS probing of candidate RNA thermometers.

T7 templates of *cspB*, *cspG*, *cspI*, *cpxP* and *lpxP*, including both 5′ UTR and CDS, were generated by PCR from DH5α gDNA using Q5® High-Fidelity 2X Master Mix (cat. M0492L, New England Biolabs). *In vitro* transcription reactions were performed using the HiScribe® T7 High Yield RNA Synthesis Kit (cat. E2040L, New England Biolabs) in 20 µl, using 1 µg of an equimolar pool of all templates. Reactions were incubated for 4 h at either 37°C or 10°C, after which RNA was probed by directly adding 200 mM final DMS to the reactions and incubating at 37°C for 2 min, or at 10°C for 30 min. Reactions were then quenched by addition of 1 volume 1 M DTT, after which RNA was cleaned up on Monarch® RNA Cleanup columns as per manufacturer instructions. Template DNA was then removed by digestion with TURBO DNase I (cat. AM2239, ThermoFisher Scientific) at 37°C for 30 min, and RNA samples were again cleaned up on Monarch® RNA Cleanup columns.

DMS-MaPseq library preparation.

DMS-MaPseq libraries were prepared as previously described[4], with minor changes. Prior to library preparation, highly abundant short RNA species, such as tRNAs, were depleted on Monarch® RNA Cleanup columns by loading a 1:1:1 mixture of total RNA in nuclease-free water, RNA binding buffer and 100% ethanol. rRNA depletion was performed on 1.1 µg total RNA using the RiboCop for Bacteria kit (cat. 126, Lexogen), with two minor changes to the manufacturer's protocol: denaturation temperature was increased to 95°C, and probe annealing temperature was lowered to 55°C. Following rRNA depletion, RNA was cleaned up on Monarch® RNA Cleanup columns, eluted in 8 µl nuclease-free water, and supplemented with 2 µl 100 µM random hexamers, 2 µl dNTPs (10 mM each) and 4 µl 5X RT Buffer [250 mM Tris-HCl pH 8.3; 375 mM KCl; 15 mM $MgCl_2$]. Samples were then incubated at 94°C for 5.5 min, to simultaneously denature and fragment the RNA to a median size of 200 nt, and immediately transferred to ice for 1 min. Samples were then supplemented with 1 µl DTT 0.1 M, 20 U SUPERase•In™ RNase Inhibitor, 200 U TGIRT™-III Enzyme (InGex, cat. TGIRT50) and 25 ng/µl actinomycin D (cat. A1410,

Merck), and then incubated at 25°C for 10 min, 57°C for 1 h, and 60°C for 1 h. Addition of actinomycin D increased strand-specificity by ~10% (*data not shown*). TGIRT-III was degraded by adding 2 µg Proteinase K and incubating at 37°C for 20 min. Proteinase K was inactivated by addition of Protease Inhibitor Cocktail (cat. P8340, Merck). cDNA-RNA hybrids were then converted to dsDNA using the NEBNext® Ultra™ II Directional RNA Second Strand Synthesis Module (cat. E7550, New England Biolabs), by incubating at 16°C for 1 h. DsDNA was cleaned up with 1.8 volumes NucleoMag NGS Clean-up and Size Select beads (cat. 744970, Macherey Nagel), and used as input for the NEBNext® Ultra™ II DNA Library Prep Kit for Illumina (cat. E7645S, New England Biolabs), as per manufacturer instructions.

Cloning of *cspG*, *cpxP* and *lpxP* constructs, and mutagenesis of *lpxP*.

Wild-type *cspG*, *cpxP* and *lpxP* FLAG-tagged, IPTG-inducible constructs, including both 5′ UTR and CDS, were prepared by amplifying the relevant regions from DH5α gDNA and by cloning them in pET22b(+) vector (cat. 69744, Merck), between the XbaI and EcoRI sites. The exact TSS was determined from DMS-MaPseq coverage. Similarly, *cspG*, *cpxP* and *lpxP* FLAG-tagged, IPTG-inducible constructs, including the sole CDS, were cloned in pET22b(+), between the NdeI and EcoRI sites. For *cpxP*, as the identified candidate thermometer encompassed part of the CDS, the CDS was cloned starting at the third in-frame ATG codon, by exploiting a naturally-occurring, in-frame NdeI site. As RNA co-transcriptional folding can be influenced by the speed of the RNA polymerase, the vector's T7 promoter was replaced with a tac promoter. The $SL_{alt}$-stabilized *lpxP* 5' UTR mutant was prepared using the Q5® Site-Directed Mutagenesis Kit (cat. E0554S, New England Biolabs), as per manufacturer instructions. All cloning steps were performed in NEB® 5-alpha Competent *E. coli* cells (cat. C2987H, New England Biolabs). All vectors were verified by Sanger sequencing (Macrogen Europe). The sequences of primers used for cloning and mutagenesis are available in Table S6. The vector containing the wild-type *lpxP* gene (inclusive of 5' UTR) has been deposited to Addgene (plasmid #212594).

Western-blot analysis of protein expression at 37°C vs. 10°C.

Sanger-verified vectors were transformed in BL21(DE3) Competent *E. coli* cells (cat. C2627H, New England Biolabs). Two independent colonies were picked and inoculated in 3 mL LB broth and grown overnight at 37°C with shaking. The next day bacteria were diluted to $OD_{600}$ ≈ 0.05 and grown until $OD_{600}$ ≈ 0.3. At this point IPTG was added to a final concentration of 1 mM, and cells were incubated with shaking at 37°C for 30 min. Bacteria were then split into two separate aliquots, pelleted, and resuspended in either 37°C or 10°C LB broth. Bacteria were then grown with shaking at either 37°C or 10°C, and 2 mL aliquots were collected after 30 min, 1 h or 2 h. Collected bacteria were pelleted and pellets were resuspended in 60 µl Lysis Buffer [10 mM Tris-HCl pH 8.0; 10 mM EDTA; 0.1% Triton X-100], supplemented with 1 µg/µl Lysozyme and 1:100 dilution Protease Inhibitor Cocktail. Samples were then subjected to 10 cycles of sonication (5 sec ON, 5 sec OFF) using a UP200St (Hielscher) ultrasonic processor. Protein concentrations were determined using Pierce™ BCA Protein Assay Kit (cat. 23225, ThermoFisher Scientific), as per manufacturer instructions. 30 µg of lysate were resolved on 10% SDS-PAGE gels, followed by transfer to nitrocellulose membrane using the iBlot™ 2 Gel Transfer system (cat. IB21001,

ThermoFisher Scientific). Membranes were blocked by incubation for 1 h in 5% (w/v) non-fat dry milk (cat. A0830, PanReac AppliChem ITW Reagents) in PBS, supplemented with 0.001% final Tween-20. Immunoblotting was performed using monoclonal ANTI-FLAG® M2 antibody (cat. F1804, Merck), or anti-LacI [9A5] universal antibody (cat. EG1501, Kerafast), and Immobilon Forte Western HRP substrate (cat. WBLUF0100, Merck).

Analysis of lpxP expression in csp knock outs.

Wild type and csp knock out BW25113 *E. coli* cells from the KEIO collection[49] were first made competent using the Mix & Go! E. coli Transformation Kit (cat. T3001, Zymo Research) and then transformed with the IPTG-inducible *lpxP* vector (see "Cloning of *cspG*, *cpxP* and *lpxP* constructs, and mutagenesis of *lpxP*" above). Two independent colonies were picked and inoculated in 3 mL LB broth and grown overnight at 37°C with shaking. The next day bacteria were diluted to $OD_{600}$ ≈ 0.05 and grown until $OD_{600}$ ≈ 0.5. At this point 1 mL of bacteria was directly mixed with 1 mL ice-cold LB broth containing 0.02 mM IPTG, and bacteria were incubated at 10°C for 1 h with moderate shaking (800 RPM). Lysis and western blot analysis were conducted as previously described (see "Western-blot analysis of protein expression at 37°C vs. 10°C" above). Knock out of csp-encoding genes was validated by PCR on gDNA from the individual clones.

*In vitro* transcription-translation using the PURE system.

*In vitro* translation analysis of full-length wild type and $SL_{alt}$-stabilized mutant, or CDS-only *lpxP*, was performed using the PURExpress® In Vitro Protein Synthesis Kit (cat. E6800S, New England Biolabs). Reactions were conducted in a final volume of 6.25 µl, using 2.5 µl Solution A, 1.875 µl Solution B, 0.1 µl SUPERase•In™ RNase Inhibitor, and ~50 fmol of pET22b(+) template (harboring a T7 promoter instead of a tac promoter). Reactions were incubated at 37°C for 1.5 h, then immediately mixed with 2X loading dye and resolved on a 12% polyacrylamide gel.

Processing of DMS-MaPseq data.

Following sequencing, paired-end reads were clipped of sequencing adapters using Cutadapt v4.4[53] (parameters: *-A AGATCGGAAG -a AGATCGGAAG -m 100:100 -O 1*), and merged using PEAR v0.9.11[54] (parameters: *-n 100 -q 20 -u 0 -e -y 10G -z*). Merged reads were then combined with R1 and the reverse-complemented R2 for read pairs that could not be merged. Next, a comprehensive annotation of *E. coli* transcriptional units with experimentally-determined TSSs was built by aggregating 5' UTR information from RegulonDB [28] and transcriptional units from EcoCyc[55], and the corresponding sequences were extracted from the *E. coli* str. K-12 substr. MG1655 genome (GenBank: U00096.3). For the analysis of known riboswitches, a reference was built including the sole riboswitch regions, ± 50 nt. Reads were then mapped to this reference using the *rf-map* tool of the RNA Framework[56] v2.8.3 and Bowtie2[57] v2.3.5.1, after clipping terminal bases with Phred quality < 20, discarding reads containing internal Ns, and trimming the 6 5'-most bases to account for possible mispriming artifacts (parameters: *-b2 -cq5 20 -ctn -cmn 0 -cl 50 -mp "--very-sensitive-local --nofw" -b5 6*). Alignments in SAM format were sorted and converted to BAM format using Samtools[58] v1.15.1. BAM alignments were then processed using RNA Framework's *rf-count* to generate both RC files (containing per-base mutations and coverage) and MM files (containing a map of mutated positions per read). Aligned reads spanning less than 100 nt of a transcript, as well as reads having more than 10% mutated bases, or less than 2 mutations, were discarded. Insertions and ambiguously aligned deletions, as well as deletions longer than 1 nt, were ignored. Mutations were considered only if both the mutated base and the two surrounding bases had Phred quality > 20, and consecutive mutations falling within 3 nt from each other were ignored (parameters: *-m -mm -wl 2000 -ds 100 -es -na -ni -md 1 -dc 3 -me 0.1 -mpr 2*). DMS-MaPseq data from total RNA, used for the calibration of folding parameters (see "Optimization of folding parameters" below), was analyzed with minor changes to the above protocol. Briefly, reads were mapped to a reference composed only of the 16S and 23S rRNA sequences. The minimum length spanned by reads was decreased to 90 nt (as total RNA DMS-MaPseq experiments were sequenced as single read 100 bp, in contrast to rRNA-depleted DMS-MaPseq experiments that were sequenced as paired-end 150 bp) and reads harboring < 2 mutations were also retained (parameters: *-m -ds 90 -es -na -ni -dc 3 -ow -me 0.1 -md 1*).

Optimization of folding parameters.

RC files from total RNA DMS-MaPseq experiments were processed using RNA Framework's *rf-norm* to obtain normalized reactivity profiles (parameters: *-sm 4 -nm 3 -rb AC -mm 1 -n 1000*). For secondary structure modeling, optimal slope (4.8) and intercept (-0.8) values were then identified via jackknifing, by simultaneously optimizing over both *in vivo* and *ex vivo* deproteinized DMS-MaPseq data from both DH5α and TOP10 cells, using RNA Framework's *rf-jackknife* (parameters: *-rp '-md 600' -x -m*), ViennaRNA v2.5.1[59], and the modified Fowlkes-Mallows index (mFMI) metric[5].

Ensemble deconvolution analysis.

Ensemble deconvolution was performed using the DRACO algorithm[4]. Briefly, DRACO slides a window of a user-defined length along each transcript, retaining only those reads falling entirely within the window's boundaries. For each window, a graph is then generated by exploiting the co-mutation information, so that, basically, each mutation in a read represents a vertex, and two bases observed to co-mutate within the same read are connected by an edge. The normalized Laplacian of the graph's adjacency matrix is then subjected to eigen decomposition and eigengap analysis to identify the number of coexisting RNA conformations making up the ensemble. This number is then used to perform a soft partitioning of the graph (graph-cut) in order to reconstruct the individual reactivity profiles of the different conformations, and their relative stoichiometries. In its original implementation, this graph-cut step involved randomly initializing the weight of each vertex for each conformation $N$ times (with $N = 50$), followed by selection of the set of weights yielding the lowest normalized graph-cut score. This initial set of weights was then iteratively altered by a factor $\varepsilon = \frac{1}{2C}$, where $C$ was the number of conformations making up the ensemble, until the normalized graph-cut score was minimized. As this procedure was performed only once, the risk was that the identified set of weights would represent only a local minimum of the graph-cut score, rather than the true minimum, potentially leading to inconsistent conformation reconstructions across consecutive DRACO runs. Furthermore, the value of $\varepsilon$ was typically too large to enable the accurate re-weighting of the vertices (for instance, with $C = 2$, $\varepsilon = 0.25$). To address these issues, we have introduced the following improvements in the DRACO algorithm (available as v1.2 from the repository https://github.com/dincarnato/draco/): 1) the number of random initializations $N$ has been increased to 500 (adjustable through the *--softClusteringInits* parameter); 2) the weight factor $\varepsilon$ has been lowered to 0.005 (adjustable through the *--softClusteringWeightModule* parameter); and 3) the entire graph-cut procedure is now repeated multiple times (adjustable through the *–softClusteringIters* parameter), to ensure convergence towards the true normalized graph-cut score minimum. Prior to running DRACO, the MM files generated by *rf-count* were pre-processed using the *filterMM* utility (available from the repository https://github.com/dincarnato/labtools) to discard reads having < 2 A/C mutated bases, and regions of extremely high coverage were randomly down-sampled to achieve a maximum per-base coverage of 500,000X. DRACO analysis was performed with a window size of 100 nt, slid in 5 nt increments, requiring a minimum base coverage of 2,000X and a minimum of 2,000 reads post-filtering to perform the eigen deconvolution, and by repeating the graph-cut procedure 30 times (parameters: *--absWinLen 100 --absWinOffset 5 --minBaseCoverage 2000 --minFilteredReads 2000 --minPermutations 10 --maxPermutations 50 --firstEigengapShift 0.95 --lookaheadEigengaps 1 --softClusteringIters 30 --softClusteringInits 500 --softClusteringWeightModule 0.005*). To determine if known riboswitches could be detected as structurally-heterogeneous, the reads from the riboswitch ± 50 nt reference were processed with DRACO with the parameter *--outputRawNClusters*. A riboswitch was considered detected if at least one window encompassing it was found to populate ≥ 2 conformations.

Comparison of DH5α/TOP10 strains and 37°C/10°C conditions.

Correlation between experiments (related to Fig. 1B, Fig. S4A and Fig. S7B) was calculated on the raw mutation frequencies of A/C bases in transcriptional units for which ≥ 50% of A/C bases had coverage ≥ 10,000X, after removing outliers (raw reactivity > 0.1). The number of conformations populated by each base in the covered transcriptome (related to Fig. 1D and Fig. 3A) was determined by parsing DRACO's JSON-formatted output files. As DRACO uses a sliding window approach, consecutive overlapping windows might be found to populate different number of conformations; in such cases, overlapping bases were assigned the highest number of conformations. Windows populating different numbers of conformations between 37°C and 10°C (related to Fig. 3B) were identified as it follows. First, windows populating 1 or 2+ conformations were extracted from DRACO's JSON-formatted output files into BED format and overlapping windows were merged using the *mergeBed* tool of BEDTools[60] v2.31.0. Any portion of the windows populating 1 conformation overlapping with the windows populating 2+ conformation was removed using BEDTools' *subtractBed*. Then, windows populating 1 or 2+ conformations common to both DH5α and TOP10 at either 37°C or 10°C were identified by intersecting the corresponding sets from both experiments, using BEDTools' *intersectBed*. Only windows populating the same number of conformations in both DH5α and TOP10 were retained. Finally, common windows populating 2+ conformations in both DH5α and TOP10 at 37°C and 1 conformation in both DH5α and TOP10 at 10°C (< ensemble heterogeneity), or vice versa (> ensemble heterogeneity), as well as regions populating the same number of conformations in both strains at both temperatures (no change), were identified by intersecting the windows set determined in the previous step using BEDTools' *intersectBed*. Window coordinates were then intersected with gene coordinates to identify which genes contained windows showing differential ensemble heterogeneity between 37°C and 10°C. For all analyses, only regions spanning at least 20 consecutive nucleotides were included. Gene ontology analysis was performed using DAVID [61].

Translation efficiency analysis.

Ribosome profiling and RNA-seq data for *E. coli* cells at 37°C, or shocked at 10°C for 10 min, were obtained from a previous study[30] (GEO dataset: GSE103421). Reads were aligned to the same transcriptome reference used for DMS-MaPseq analysis, using RNA Framework's *rf-map* and Bowtie [62] v1.3.1, allowing a maximum of 2 mapping positions (parameters: *-ca3 CTGTAGGCACCATCAA -bnr -ow -bm 2 -bc 32000 -ba*). After discarding all reads mapping to the rRNA operons, read counts for protein-coding genes containing (or not) windows showing differential heterogeneity between 37°C and 10°C (see "Comparison of DH5α/TOP10 strains and 37°C/10°C conditions" above) were calculated by intersecting gene coordinates (for protein-coding genes with CDS ≥ 99 bp) in BED format with the relevant BAM files, using BEDTools' *intersectBed* (parameter: *-c*). Only windows ≥ 20 nt were considered. For both Ribo-seq and RNA-seq data, per-gene RPKMs were calculated as:

$$RPKM = \frac{C}{NL} \times 1{,}000{,}000$$

where *C* was the read count on the gene, *N* was the total number of reads mapped in the experiment, and *L* was the length of the gene in kilobases. Translation efficiency for each gene (related to Fig. 3e) was then calculated as:

$$TE = \frac{RPKM_{Ribo-seq} + 0.1}{RPKM_{RNA-seq} + 0.1}$$

where 0.1 is a pseudo count added to avoid division by zero. Only genes expressed ≥ 1 RPKM both at 37°C and 10°C were considered.

Comparison of regions populating 1 vs. 2+ conformations.

Eight features were evaluated for regions populating 1 vs. 2+ conformations at 37°C, namely: AC % content, GC% content, median Shannon entropy, median unpaired probability, median reactivity, Gini index, and Z-score (related to Fig. 2A-E, S3 and S5A-E). Only regions ≥ 20 nt were considered. First, bulk reactivity profiles for both DH5α and TOP10 grown at 37°C were obtained by normalizing the respective RC files (see "Processing of DMS-MaPseq data" above) using RNA Framework's *rf-norm* (parameters: *-sm 4 -nm 3 -rb AC -mm 1 -n 1000*), and the resulting normalized XML reactivity files were combined using RNA Framework's *rf-combine*. From these XML files, reactivity data for regions populating either 1 or 2+ conformations was extracted and used to calculate the median reactivity and Gini index distributions. Combined XML files were then passed to RNA Framework's *rf-fold* to compute base-paring probabilities and Shannon entropies (parameters: *-sl 4.8 -in -0.8 -md 600 -dp -sh*). Unpaired probabilities per base were calculated as:

$$1 - \sum_{j=i}^{J} p(i,j)$$

where *p(i,j)* is the base-pairing probability between nucleotides *i* and *j*, over all possible *J* partners. For unconstrained predictions, the same parameters were used with the addition of the *-i* parameter to ignore experimental reactivities. Distributions of folding free energy Z-scores were calculated on the nucleotide sequences corresponding to the regions populating either 1 or 2+ conformations, in the absence of any constraint, using ViennaRNA. As regions populating 1 conformation were on average longer than those populating 2+ conformations, an equal number of random portions were extracted from regions populating 1 conformation to match the size distribution of the regions populating 2+ conformations. For Z-score calculation, the sequence of each region was shuffled 100 times, by preserving dinucleotide frequencies, and the corresponding folding free energies were predicted using RNAfold. The Z-score for each region was then calculated as:

$$Z = \frac{\Delta G - \mu}{\sigma}$$

where $\Delta G$ was the folding free energy for the original sequence, while $\mu$ and $\sigma$ were respectively the average and the standard deviation of the folding free energies across the 100 shuffled sequences.

Sequence-level conservation analysis.

To evaluate the conservation of regions populating 1 vs. 2+ conformations, a multiple sequence alignment was computed using Mugsy v1.2.3 [63] and the following 10 Gram-negative bacteria genomes: *Escherichia coli* str. K-12 substr. MG1655 (GenBank: U00096.3), *Salmonella enterica* subsp. enterica serovar Typhimurium str. LT2 (GenBank: AE006468.2), *Shigella flexneri* 2a str. 2457T (GenBank: AE014073.1), *Klebsiella pneumoniae* subsp. pneumoniae HS11286 (GenBank: CP003200.1), *Yersinia pestis* CO92 (GenBank: AL590842.1), *Enterobacter* sp. 638 (GenBank: CP000653.1), *Serratia marcescens* strain KS10 (GenBank: CP027798.1), *Pectobacterium carotovorum* strain WPP14 (GenBank: CP027798.1), *Shigella dysenteriae* strain SWHEFF_49 (GenBank: CP055055.1), and *Enterobacter cloacae* isolate 1382 (GenBank: OW968328.1). The resulting alignment was parsed to calculate the % conservation at each position, with respect to the *E. coli* genome.

Reactivity profile reconstruction and structure modeling for high-confidence regions.

High-confidence structurally-heterogeneous regions for which the deconvolved reactivity profiles could be non-ambiguously matched between DH5α and TOP10 (average correlation of reactivity profiles ≥ 0.65) were extracted using RNA Framework's *rf-json2rc*, by including 20 extra bases on either side of the structure (parameters: *-ec 1000 -mom 0.35 -e 20 -cf 0.1 -i 0.1 -mcm 0.65 -mcr 0.65*). The tool processes DRACO's JSON-formatted output files from two experiments, aggregating those regions showing sufficient agreement between the deconvolved reactivity profiles across the two experiments, yielding two RC files containing the per-base coverage and mutations across the different conformations reconstructed by DRACO for the analyzed RNAs. The resulting RC files were then processed using RNA Framework's *rf-norm* to yield normalized reactivity profiles (parameters: *-sm 4 -nm 3 -rb AC -mm 1 -n 100*). Structure modeling was performed using the *consensusFold* utility (available from the repository https://github.com/dincarnato/labtools), which leverages RNAalifold[64] to aggregate multiple reactivity profiles into a consensus secondary structure (parameters: *-sl 4.8 -in -0.8 -md 600*). For the modeling of secondary structures under cold shock conditions an additional parameter *-t 10* was specified to set the folding temperature to 10°C.

Covariation analysis.

To evaluate the conservation of the identified structures, we implemented the evolutionary conservation analysis module of the DeConStruct framework, built on top of the *cm-builder* pipeline (available from the repository https://github.com/dincarnato/labtools) we previously introduced[4,33] (which exploits Infernal v1.1.3[65] and R-scape v2.0.0.q[35,66]) to be able to handle full bacterial genomes rather than just individual transcripts. For each predicted structure (filtering out those with a known match in Rfam[67]) a covariance model (CM) was first built using Infernal's *cmbuild* and the sole *E. coli* sequence. The CM was then used to search a database of 7598 representative archeal and bacterial genomes (and associated plasmids, when present) from RefSeq to iteratively identify putative homologs. In its original implementation, *cm-builder* used an E-value-based approach to search in the database. This approach had two main limitations. First, the E-value for the identified matches was dependent on the size of the searched database, potentially leading to different results with different database sizes. Second, it required the calibration of the CMs via Infernal's *cmcalibrate* module, a computationally intensive task, not easily scalable to hundreds of candidates. To address these issues, we implemented a bit score-based search. Briefly, to trick Infernal into thinking that a CM had been calibrated, a fake set of ECMLC/ECMGC/ECMLI/ECMGI field values was introduced into the CM. These fields are only used to determine the E-value of a database search, but they do not affect the bit score. Then, a decoy database was built by randomly extracting and reversing ~10% of the sequences from the original genome database. Infernal's *cmsearch* was then used to search the CM against the decoy database. A noise threshold $N$ was defined by taking the highest possible bit score returned by this search, and by rounding it up to the nearest multiple of 5. If $N < 20$, then $N$ was set to 20. The search was then repeated against the original database, retaining only those matches having bit score > $N$. Matches having < 50% canonical base-pairs, as well as truncated hits covering < 75% of the structure were discarded. The resulting set of candidate homologs was then realigned against the original CM by using Infernal's *cmalign*. The whole procedure was repeated a maximum of 3 times. At each iteration, $N$ was increased by 10, and the alignment of candidate homologs was analyzed using R-scape's APC-corrected G-test statistics and a relaxed E-value threshold of 0.1 (to account for those structures falling within coding regions for which sequence variation might be "constrained" by the underlying amino acid sequence). In case the number of significantly covarying base-pairs dropped with respect to the previous iteration (except for the first iteration), the procedure was stopped. The final alignment was then polished by discarding sequences having a length that was significantly different from the majority of the sequences in the alignment. This was achieved by converting sequence lengths to z-scores, and by discarding sequences with abs(z-score) > 2 and > 10% length difference with respect to the average sequence length in the alignment (implemented in the *stockholmPolish* tool available from the repository https://github.com/dincarnato/labtools). To further select only high-confidence alignments, we performed a stringent filtering by selecting alignments matching the following 3 criteria: 1) ≥ 25% of the helices showing helix-level covariation (R-scape's Lancaster aggregated E-value < 0.05); 2) ≥ 12.5% of the base-pairs showing covariation (R-scape APC-corrected G-test statistics E-value < 0.1); and 3) ≥ 5 base-pairs showing covariation.

<u>Evaluation of energy barriers and fraction changed base-pairs between conformations.</u>

Transition barriers were estimated on the set of structures predicted from structurally-heterogeneous regions whose DRACO-deconvolved reactivity profiles could be non-ambiguously matched between DH5α and TOP10 cells (see "Reactivity profile reconstruction and structure modeling for high-confidence regions" above). Estimation was performed using DrFindpath, a component of the DrTransformer package [68]. DrFindpath employs the Findpath heuristic [69], which is implemented in the Vienna RNA library. The fraction of changed base-pairs between conformations was calculated as:

$$F = 1 - \frac{C}{c_1 + c_2 + C}$$

where $C$ was the number of base-pairs common to both conformations, and $c_1$ and $c_2$ were respectively the numbers of base-pairs unique to either conformations.

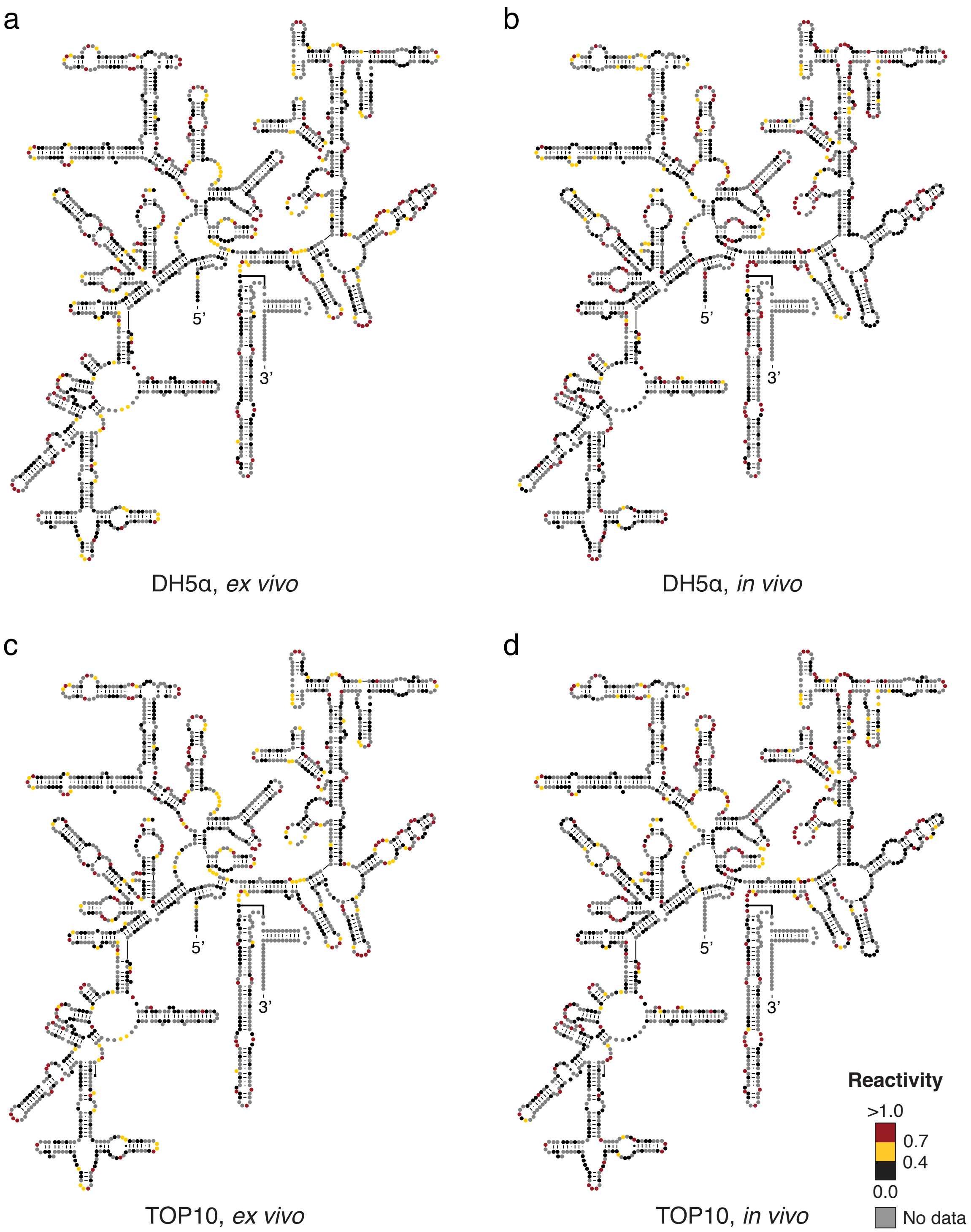


**Supplementary Figure 1.**

DMS reactivities overlaid onto the phylogenetically-accepted *E. coli* 16S rRNA structure. (**a**) DH5α, *ex vivo* deproteinized. (**b**) DH5α, *in vivo*. (**c**) TOP10, *ex vivo* deproteinized. (**d**) TOP10, *in vivo*.

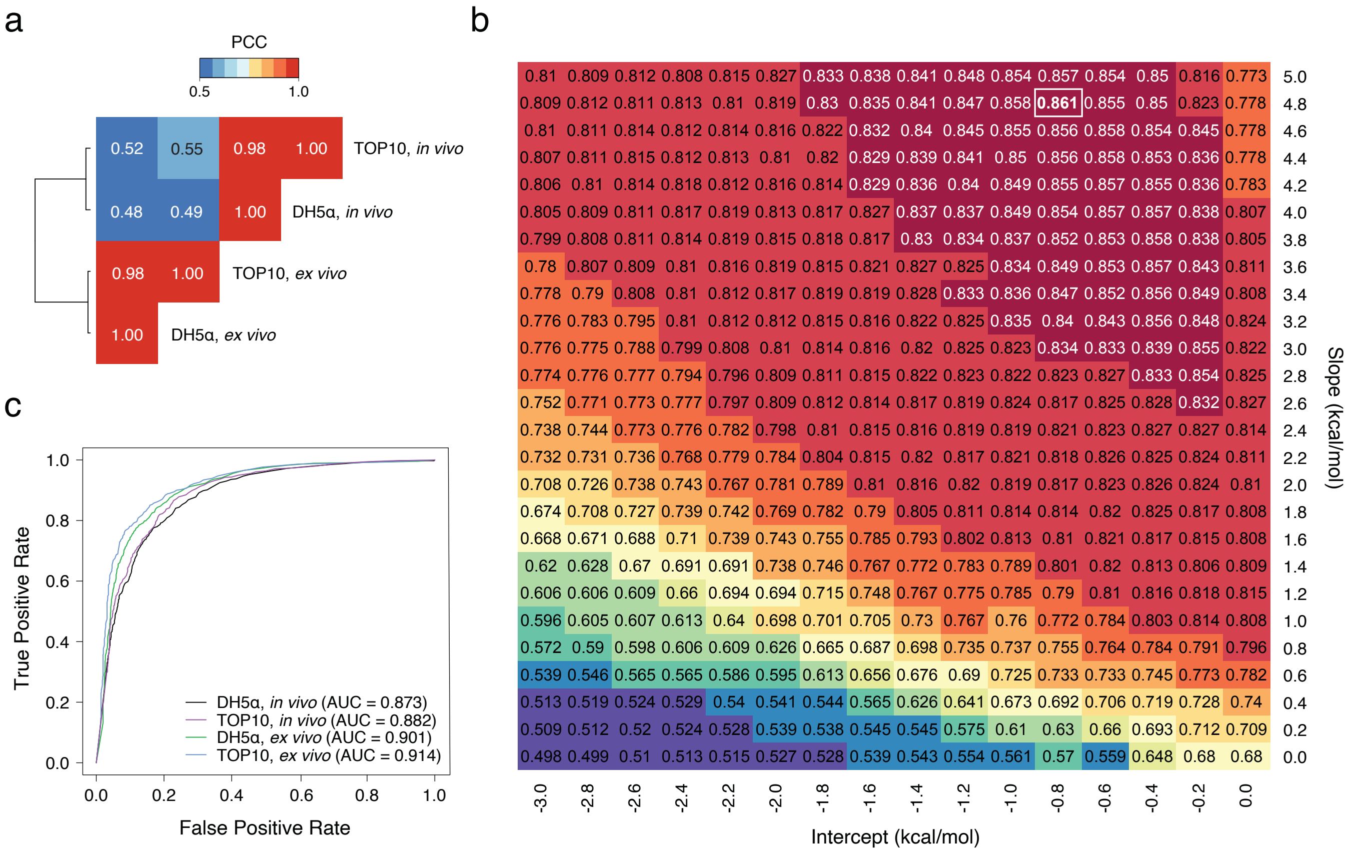


**Supplementary Figure 2.**

(**a**) Heatmap of pairwise Pearson correlation coefficients (PCC) of raw DMS reactivities across the 16S and 23S rRNAs of DH5α and TOP10 cells, probed either *in vivo* or *ex vivo* post-deproteinization. (**b**) Heatmap of mFMI values as determined by jackknifing of optimal slope and intercept parameters on 16S and 23S rRNAs. (**c**) Received Operator Characteristic (ROC) curves for DMS reactivities across the 16S and 23S rRNAs of DH5α and TOP10 cells, probed either *in vivo* or *ex vivo* post-deproteinization.

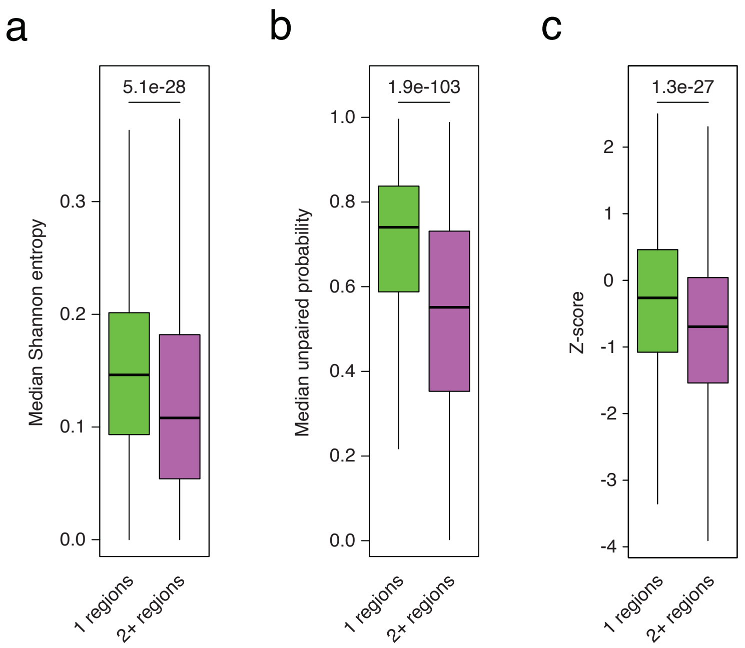


**Supplementary Figure 3.**

Box-plots depicting the distributions for different features across regions populating 1 or 2+ conformations *in vivo*. (**a**) Median Shannon entropies (from DMS-constrained predictions). (**b**) Median unpaired probabilities (from DMS-constrained predictions). (**c**) Z-scores of folding free energies. For all plots, boxes span the 25$^{th}$ to the 75$^{th}$ percentile. The center represents the median. Outliers (values below the 25$^{th}$ percentile – 1.5 times the IQR, or above the 75$^{th}$ percentile + 1.5 times the IQR) are not shown. P-values are calculated using the Wilcoxon rank sum test.

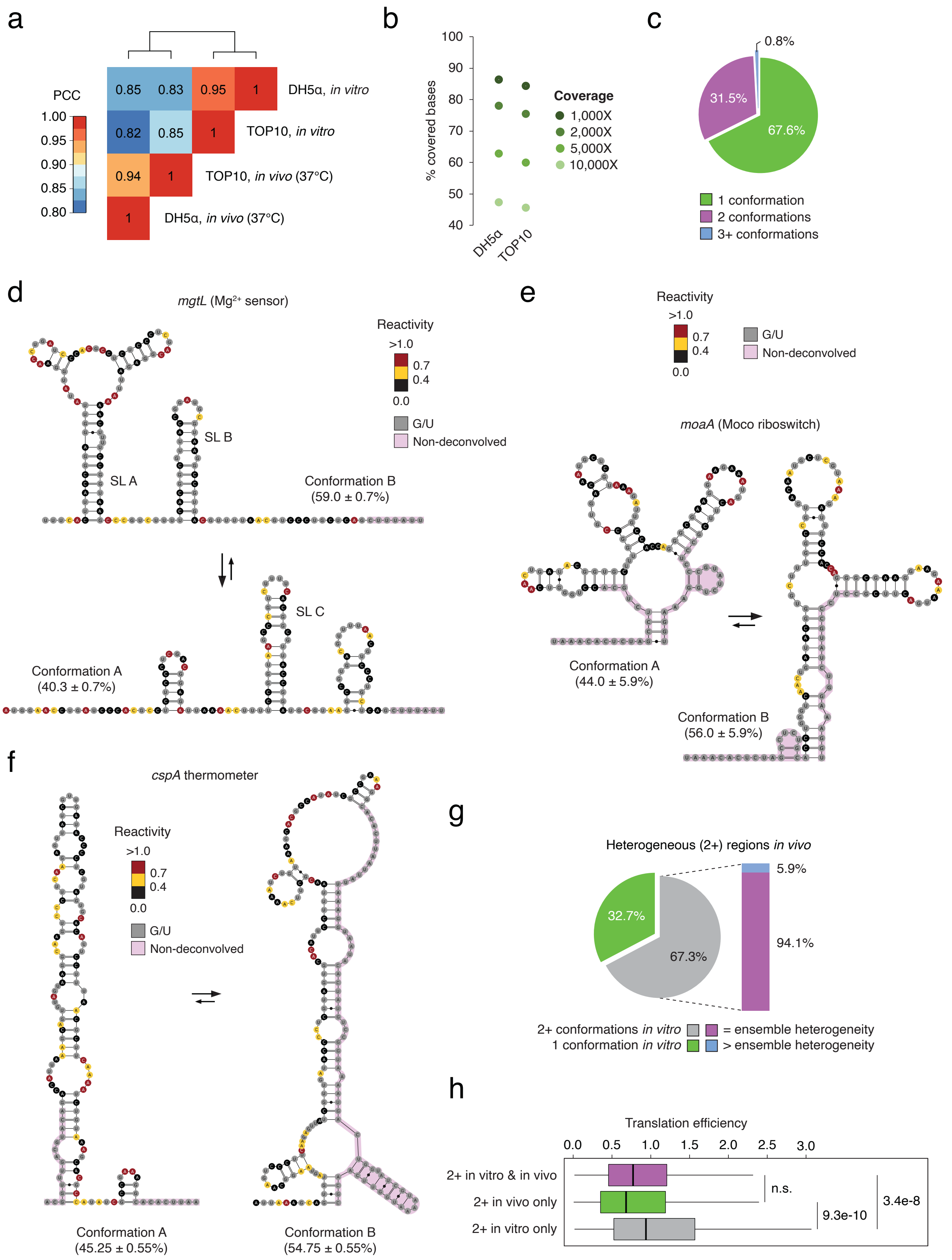


*(legend on the next page)*

**Supplementary Figure 4.**

(**a**) Heatmap of pairwise Pearson correlation coefficients (PCC) of raw DMS reactivities across bases with coverage ≥ 10,000X in the transcriptome of DH5α and TOP10 cells, for *in vivo* and *in vitro* refolded conditions. Outliers (bases with mutation frequency > 0.1) were excluded. (**b**) Percentage of bases covered in the expressed transcriptome (TPM ≥ 10) of DH5α and TOP10 cells, at different sequencing depths, for the *in vitro* refolded samples. (**c**) Pie-chart depicting the percentages of bases in the *E. coli* transcriptome populating 1, 2, or 3+ conformations *in vitro*. Only bases populating the same number of conformations in DH5α and TOP10 cells were considered. (**d**) Secondary structure of the two known conformations of the *mgtL* $Mg^{2+}$ sensor, with overlaid DMS reactivities, averaged across DH5α and TOP10 *in vitro* refolded samples. Bases falling outside of the region deconvolved by DRACO are marked in pink. (**e**) Secondary structure of the two conformations of the *moaA* Moco riboswitch, with overlaid DMS reactivities, averaged across DH5α and TOP10 *in vitro* refolded samples. Conformation A corresponds to the known structure of the riboswitch, while conformation B has been derived by DMS-constrained structure modelling. (**f**) Secondary structure of the two known conformations of the *cspA* thermometer, with overlaid DMS reactivities, averaged across DH5α and TOP10 *in vitro* refolded samples. (**g**) Pie-chart depicting the fraction of 2+ regions identified *in vivo*, populating 1 or 2+ conformations *in vitro*. The bar inset further displays the fraction of regions showing structural heterogeneity both *in vivo* and *in vitro* that populate the same, or an increased number of conformations, *in vitro* as compared to *in vivo*. (**h**) Box-plot depicting the translation efficiency calculated on regions populating 2+ conformations both *in vitro* and *in vivo*, only *in vivo*, or only *in vitro*. Boxes span the $25^{th}$ to the $75^{th}$ percentile. The center represents the median. Outliers (values below the $25^{th}$ percentile – 1.5 times the IQR, or above the $75^{th}$ percentile + 1.5 times the IQR) are not shown. P-values are calculated using the Wilcoxon rank sum test.

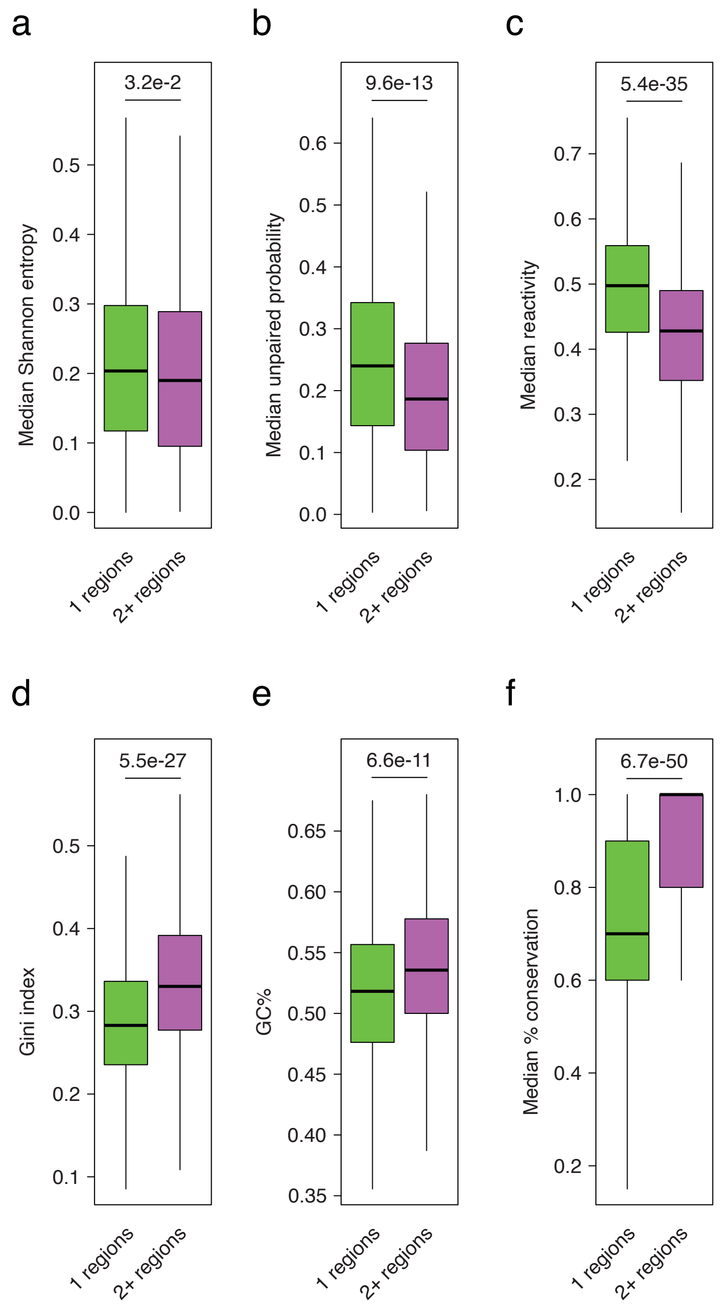


**Supplementary Figure 5.**
Box-plots depicting the distributions for different features across high-confidence regions (regions common to both *in vitro* and *in vivo* experiments, subtracted of RNase E sites) populating 1 or 2+ conformations. (**a**) Median Shannon entropies (from unconstrained predictions). (**b**) Median unpaired probabilities (from unconstrained predictions). (**c**) Median bulk *in vivo* DMS reactivities. (**d**) Gini indexes calculated on bulk *in vivo* DMS reactivities. (**e**) GC% content. (**f**) Median % sequence conservation calculated on a set of 10 Gram-negative bacterial genomes. For all plots, boxes span the 25$^{th}$ to the 75$^{th}$ percentile. The center represents the median. Outliers (values below the 25$^{th}$ percentile – 1.5 times the IQR, or above the 75$^{th}$ percentile + 1.5 times the IQR) are not shown. P-values are calculated using the Wilcoxon rank sum test.

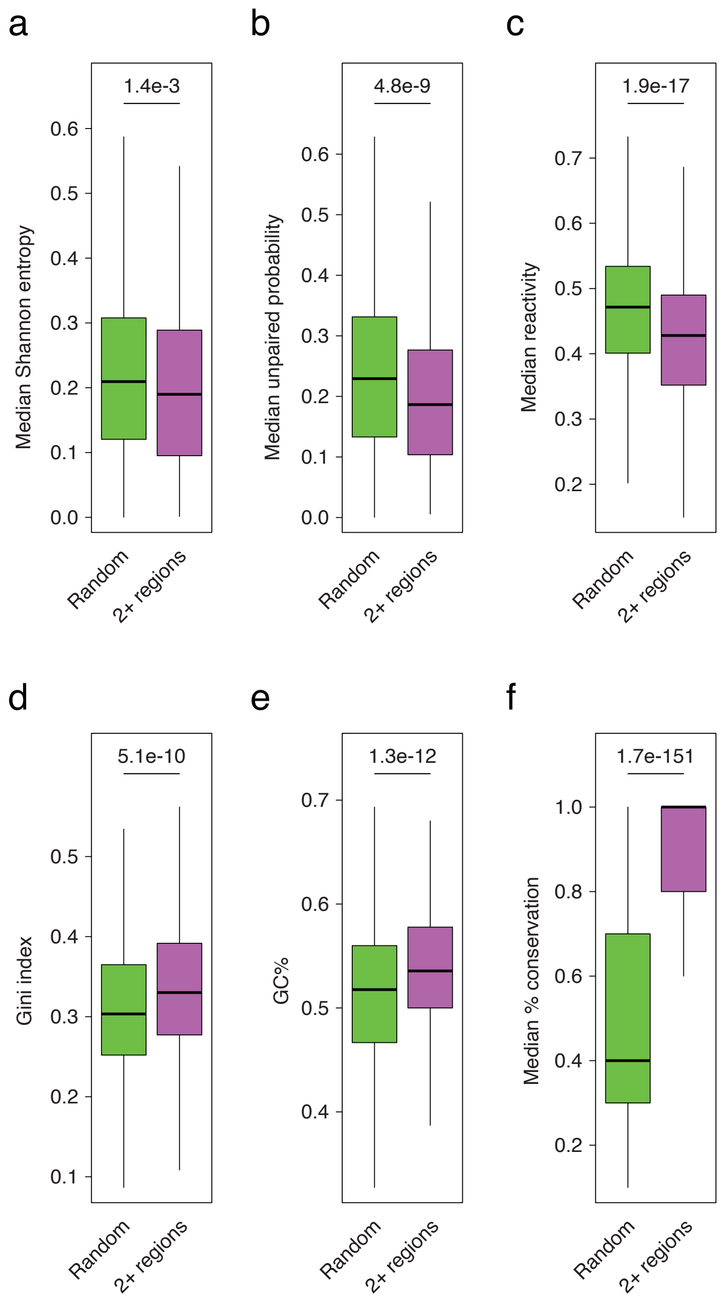


**Supplementary Figure 6.**
Box-plots depicting the distributions for different features across high-confidence regions (regions common to both *in vitro* and *in vivo* experiments, subtracted of RNase E sites) populating 2+ conformations, as compared to random transcriptome regions of matching size. (**a**) Median Shannon entropies (from unconstrained predictions). (**b**) Median unpaired probabilities (from unconstrained predictions). (**c**) Median bulk *in vivo* DMS reactivities. (**d**) Gini indexes calculated on bulk *in vivo* DMS reactivities. (**e**) GC% content. (**f**) Median % sequence conservation calculated on a set of 10 Gram-negative bacterial genomes. For all plots, boxes span the 25$^{th}$ to the 75$^{th}$ percentile. The center represents the median. Outliers (values below the 25$^{th}$ percentile – 1.5 times the IQR, or above the 75$^{th}$ percentile + 1.5 times the IQR) are not shown. P-values are calculated using the Wilcoxon rank sum test.

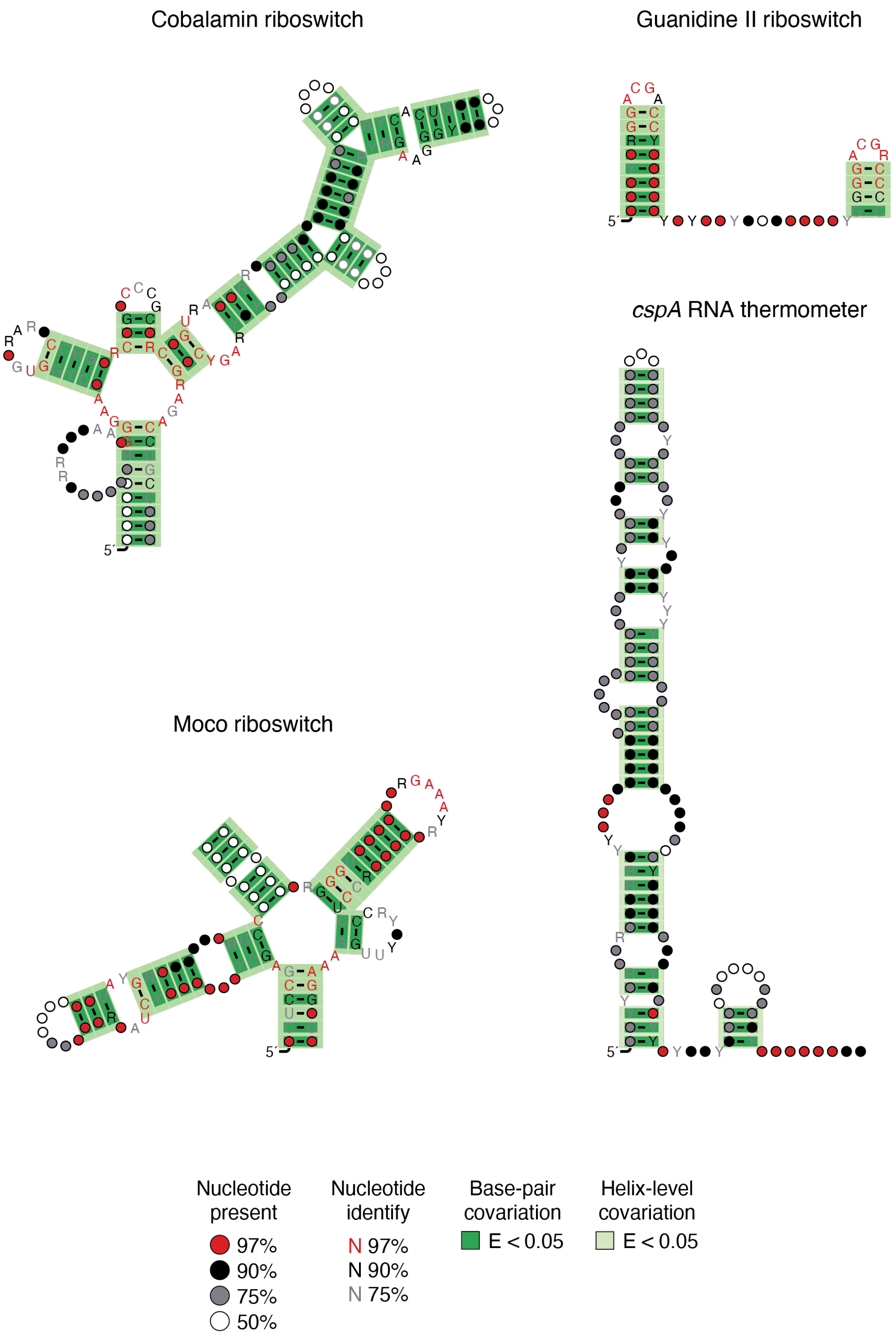


**Supplementary Figure 7.**

Validation of the covariation analysis module of the DeConStruct pipeline. Sample structure models for known bacterial RNA switches, as inferred by automatic phylogenetic analysis. Base-pairs showing significant covariation (as determined by R-scape) are boxed in dark green (E-value < 0.05). Helices showing helix-level covariation support (E-value < 0.05) are boxed in light green.

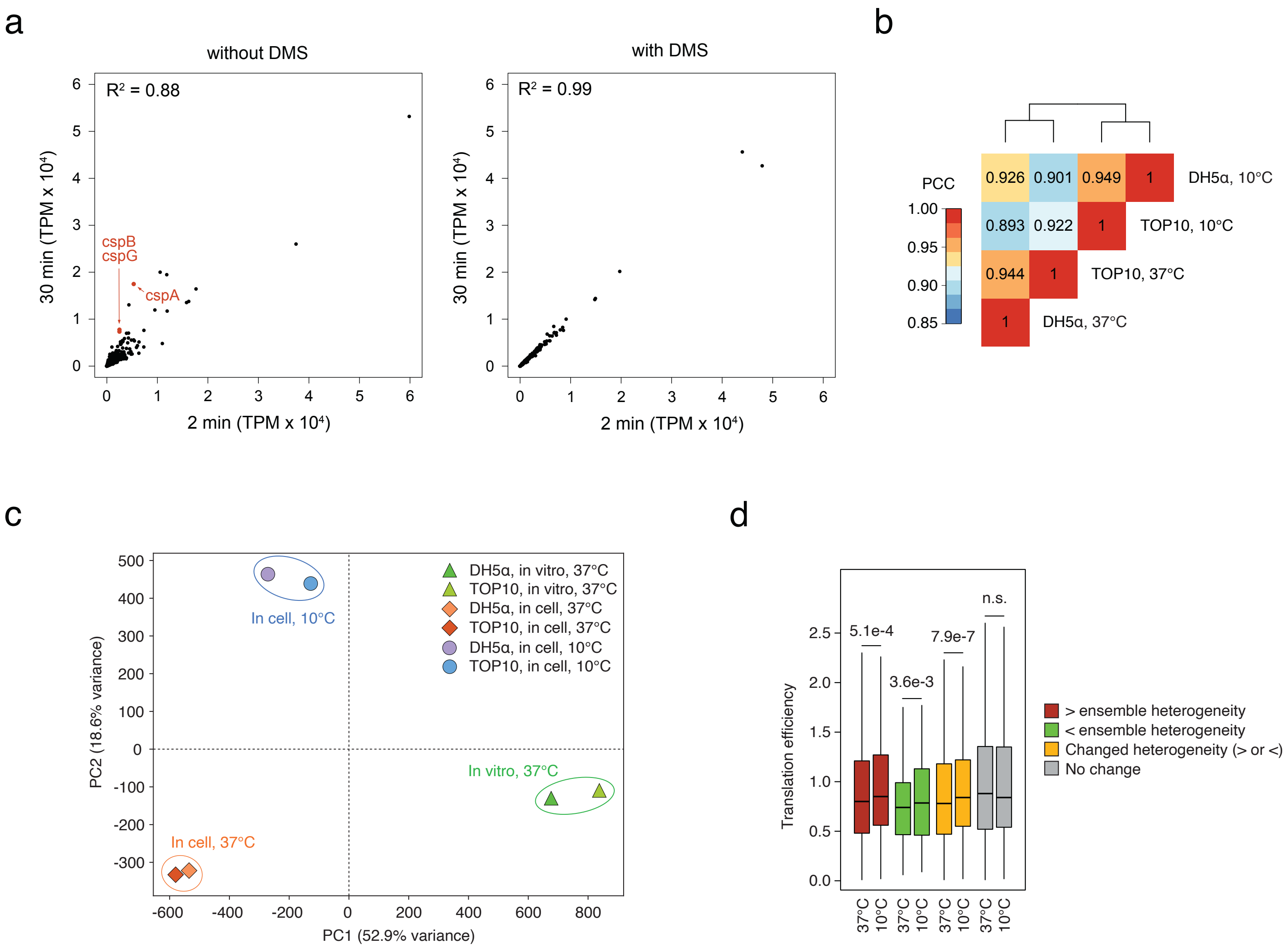


**Supplementary Figure 8.**

(**a**) Scatter plots comparing gene expression in *E. coli* cells subjected to 20 min cold shock, and then treated either with 200 mM DMS or ethanol (vehicle) for 2 or 30 min. (**b**) Heatmap of pairwise Pearson correlation coefficients (PCC) of raw DMS reactivities across bases with coverage ≥ 10,000X in the transcriptome of DH5α and TOP10 cells, for *in vivo* 37°C and 10°C conditions. Outliers (bases with mutation frequency > 0.1) were excluded. (**c**) Principal component analysis (PCA) performed on DMS reactivities. (**d**) Box-plot depicting translation efficiencies at 37°C vs. 10°C (10 min) calculated on protein-coding genes encompassing regions showing increased (red), decreased (green), changed (either increased or decreased, yellow) or unchanged (grey) structural heterogeneity upon cold shock as compared to 37°C. Ribosome profiling data from (*15*). Boxes span the 25$^{th}$ to the 75$^{th}$ percentile. The center represents the median. Outliers (values below the 25$^{th}$ percentile – 1.5 times the IQR, or above the 75$^{th}$ percentile + 1.5 times the IQR) are not shown. P-values are calculated using the paired Wilcoxon rank sum test.

a

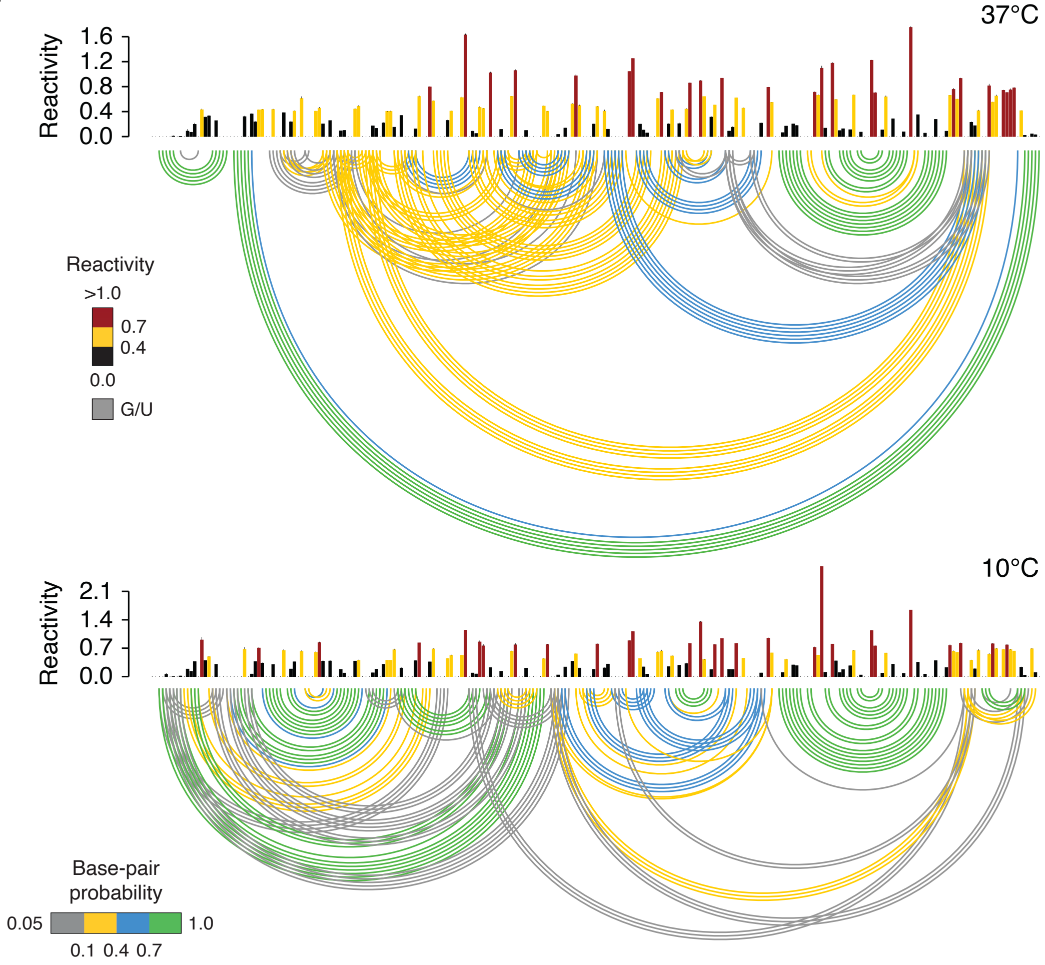


b

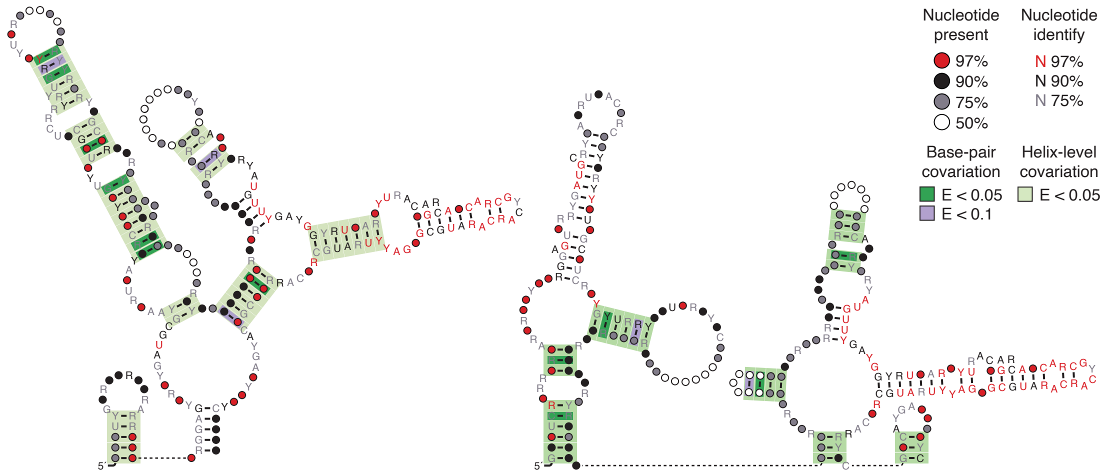


*(legend on the next page)*

**Supplementary Figure 9.**

(**a**) Reactivity profiles and base-pairing probabilities for *cpxP*'s 5′ UTR at 37°C and 10°C. Reactivities are averaged across two independent experiments. Error bars represent the standard deviation. (**b**) Structure models for the two conformations of the identified *cpxP* RNA thermometer, inferred by phylogenetic analysis. Base-pairs showing significant covariation (as determined by R-scape) are boxed in dark green (E-value < 0.05) or purple (E-value < 0.1). Helices showing helix-level covariation support (E-value < 0.05) are boxed in light green.

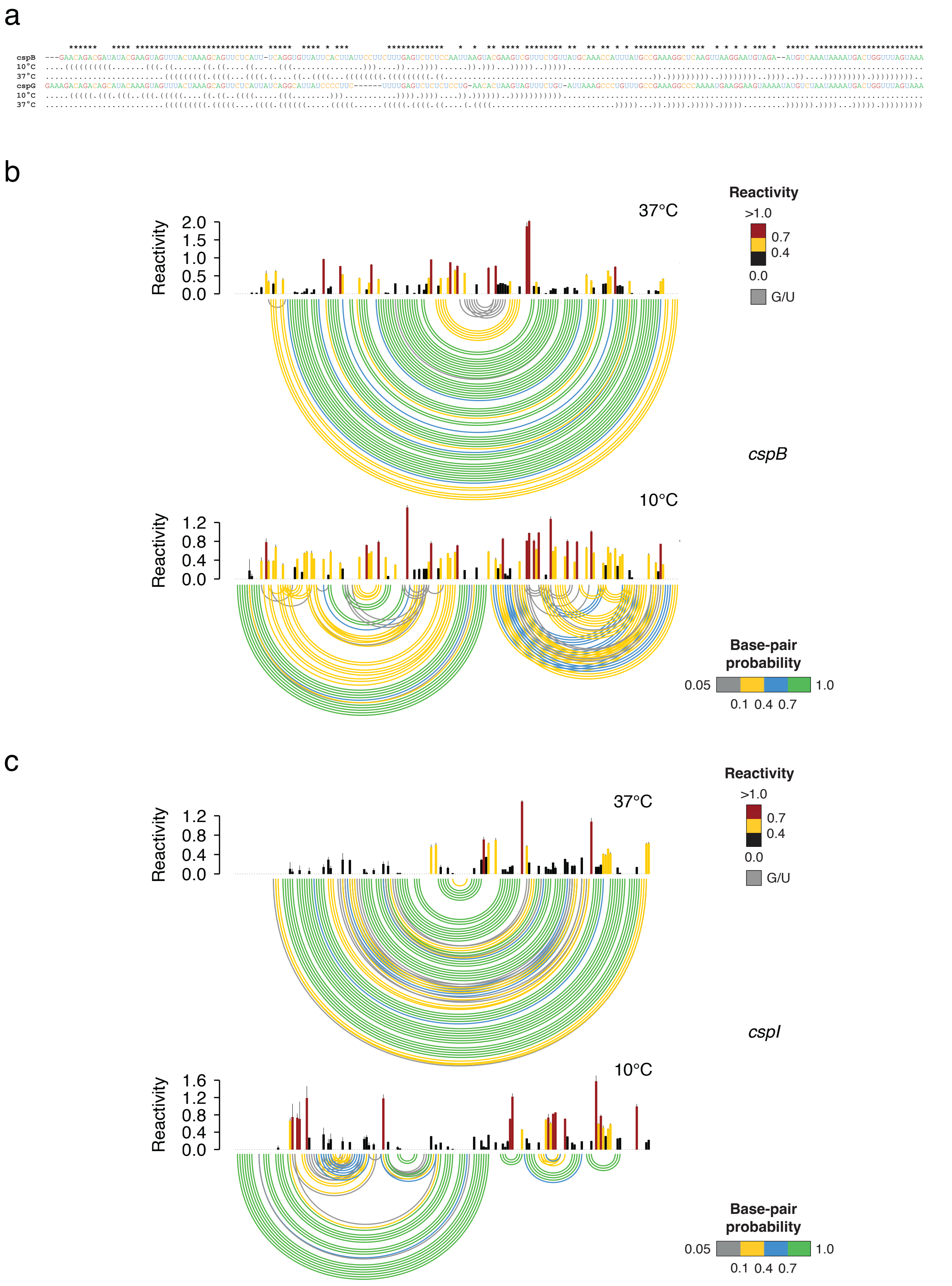


*(legend on the next page)*

**Supplementary Figure 10.**

(**a**) Sequence alignment between the 5′ UTRs of *cspG* and *cspI* (computed with Clustal Omega). Secondary structures inferred from DMS reactivities are shown in dot bracket notation. (**b**) Reactivity profiles and base-pairing probabilities for *cspB*'s 5′ UTR at 37°C and 10°C. Reactivities are averaged across two independent experiments. Error bars represent the standard deviation. (**c**) Reactivity profiles and base-pairing probabilities for *cspI*'s 5′ UTR at 37°C and 10°C. Reactivities are averaged across two independent experiments. Error bars represent the standard deviation.

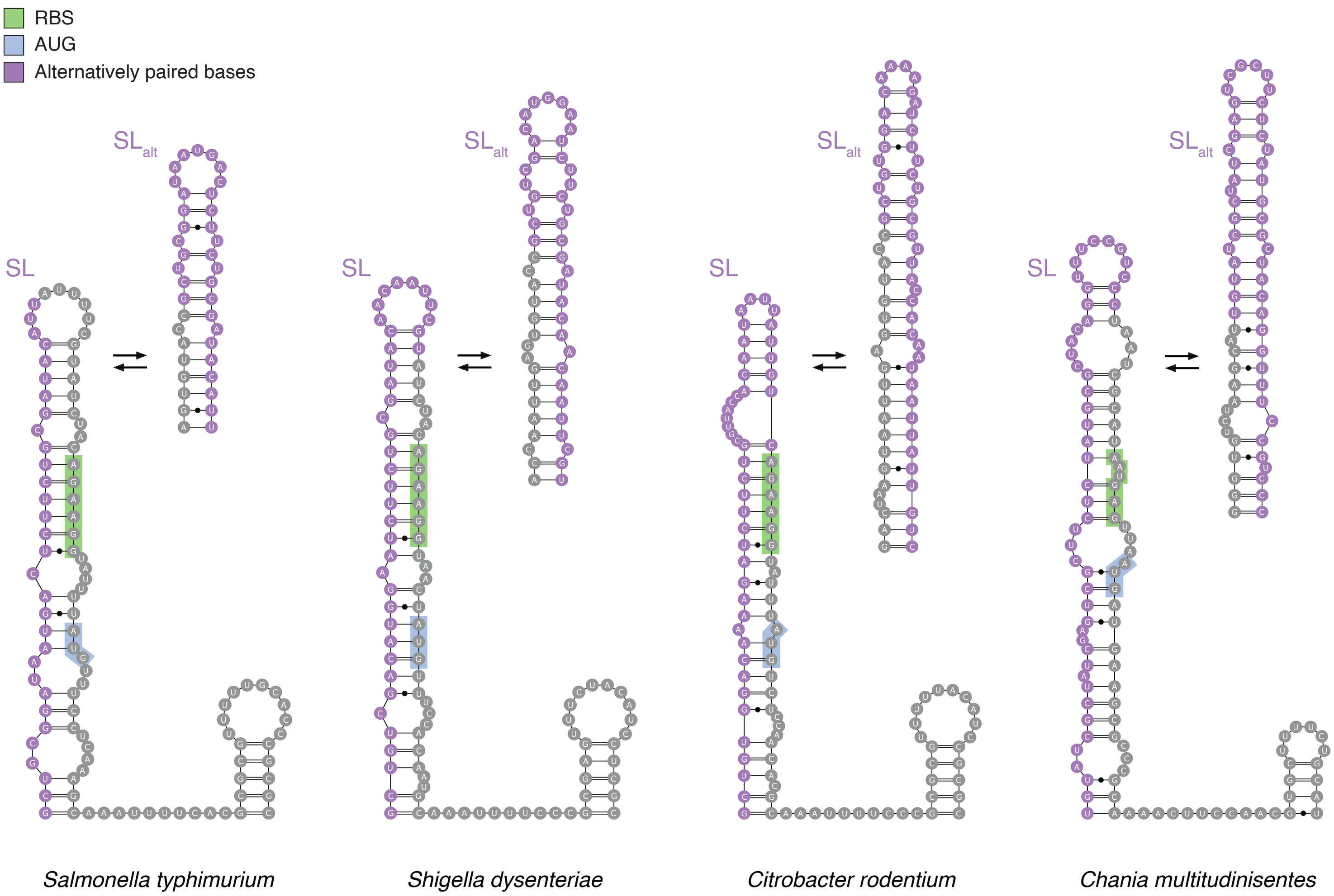


**Supplementary Figure 11.**

Examples of putative *lpxP* homologs identified by covariance model-guided search. The translation-incompetent conformation forming the SL stem-loop, which sequesters the RBS (green) and start codon (blue), is shown, along with the region undergoing structural rearrangement to form $SL_{alt}$ (purple).

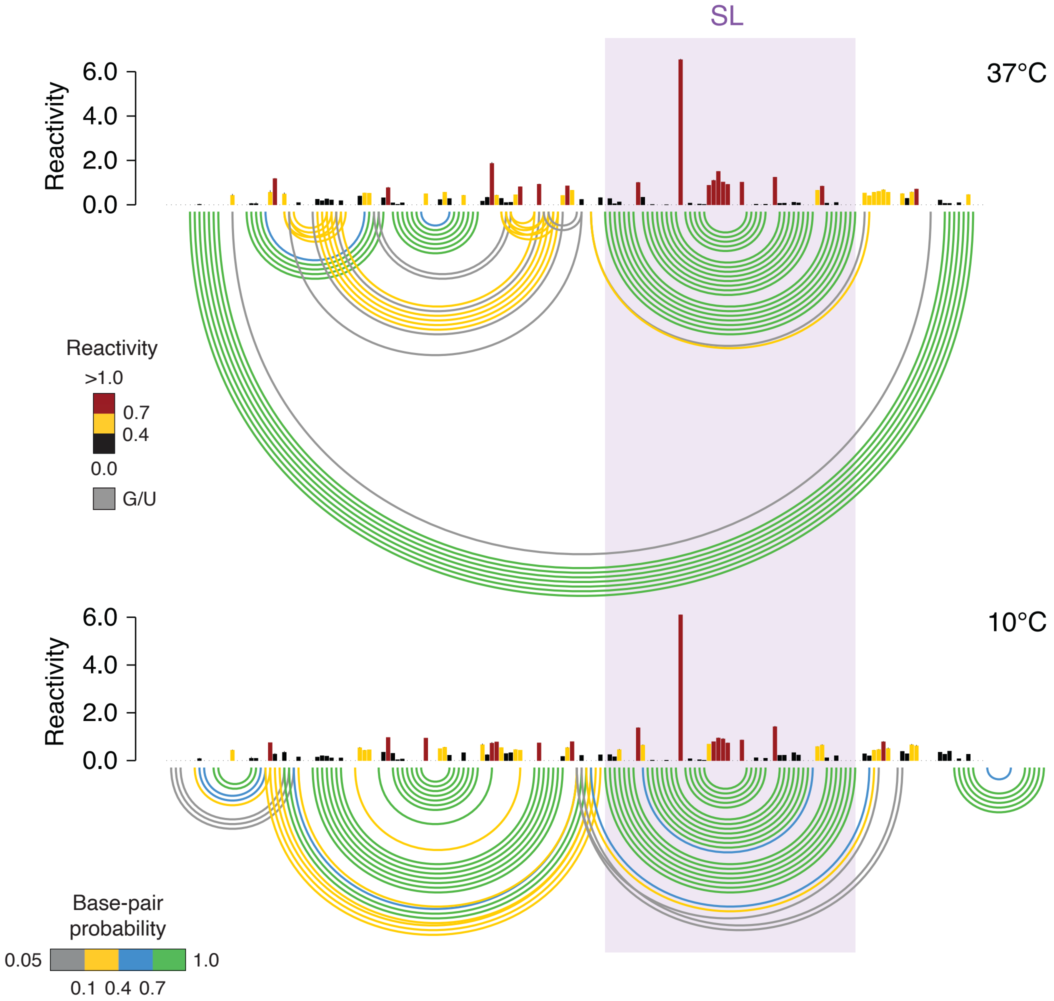


**Supplementary Figure 12.**

Reactivity profiles and base-pairing probabilities for *lpxP*'s 5′ UTR at 37°C and 10°C. Reactivities are averaged across two independent experiments. Error bars represent the standard deviation. The SL is consistently observed both at 37°C and 10°C.

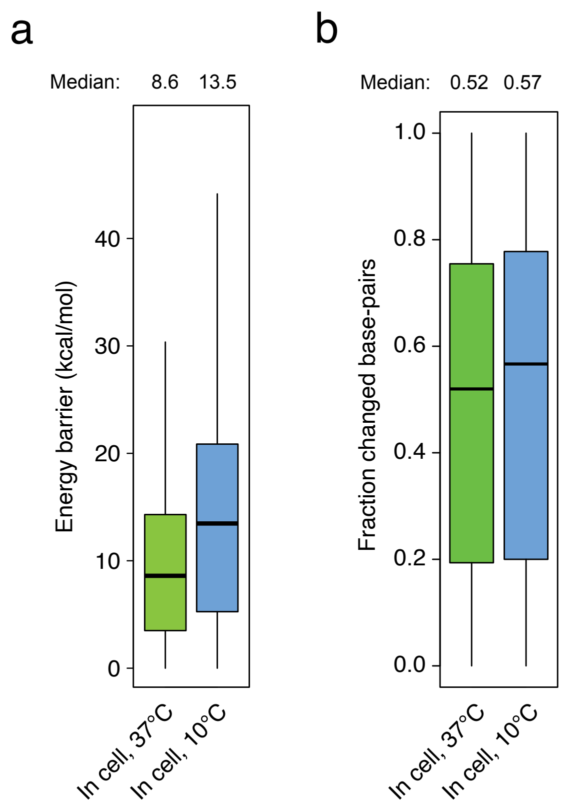


**Supplementary Figure 13.**

(**a**) Box-plot depicting the distribution of energies for the transition barriers between alternative conformations identified at 37°C and 10°C. (**b**) Box-plot depicting the faction of changed base-pairs between alternative conformations identified at 37°C and 10°C. Boxes span the 25$^{th}$ to the 75$^{th}$ percentile. The center represents the median. Outliers (values below the 25$^{th}$ percentile – 1.5 times the IQR, or above the 75$^{th}$ percentile + 1.5 times the IQR) are not shown.
